# Mapping Connectional Differences between Humans and Macaques in the Nucleus Accumbens Shell-Core Architecture

**DOI:** 10.1101/2020.06.12.147546

**Authors:** Xiaoluan Xia, Lingzhong Fan, Chen Cheng, Luqi Cheng, Long Cao, Bin He, Junjie Chen, Haifang Li, Tianzi Jiang

**Author notes:** Co-correspondence author Tianzi Jiang, Brainnetome Center, Institute of Automation, Chinese Academy of Sciences, Beijing 100190, China. Or Haifang Li, College of Computer Science and Technology, Taiyuan University of Technology, Taiyuan 030600, China. Co-first author (Xiaoluan Xia, Lingzhong Fan).

## Abstract

Two nucleus accumbens subregions, the shell and core, differ in the patterns whereby they integrate signals from prefrontal and limbic areas of the brain. In this study, we investigated whether the disproportionate volumetric differences of these brain areas, particularly the prefrontal cortex, between humans and macaques are accompanied by unique modifications of their macroscopic integrative connections with the shell and core. More specifically, we characterized the tractographic connectivity profiles of the human and macaque shell-core architecture and compared them between the two species. To make the cross-species comparisons more viable, we used the same whole-brain voxel-wise tractography-defined shell-like and core-like divisions in the two species as seeds and delineated pairs of interspecies connectionally comparable (ICC) target regions based on the similarity of the resting-state functional connectivity profiles for the two species, and finally used these seeds and ICC targets to establish a fingerprint-based common space for cross-species comparisons. Our results revealed that dissimilar structural connectivity profiles were found in the prefrontal but not the subcortical target group. We further localized this difference to specific targets to infer possible functional modifications between the two species.

## Introduction

The nucleus accumbens (Acb) is believed to integrate mnemonic and emotional signals from the prefrontal and limbic regions, helping organisms to achieve motivationally relevant goals by facilitating action selection, e.g., procuring things worth having or avoiding aversive consequences. Furthermore, in part because of their distinct structural and functional connectivity profiles, the well-documented Acb subregions, the shell and core, appear to promote distinct patterns of behavior during action selection (Floresco, 2015). Specifically, the shell receives prominent glutaminergic projections from the ventromedial prefrontal cortex (PFC), subiculum, CA1 fields of the hippocampus (HIPP), and parvicellular basolateral amygdala (AMYG) as well as small dopaminergic projections from the ventral tegmental area and functionally plays a role in suppressing low- or non-rewards stimuli that may interfere with the best available reward-predicting stimuli utilizing value-driven decision making (Groenewegen et al., 1987; Wright and Groenewegen, 1996; Heidbreder and Groenewegen, 2003; Stopper and Floresco, 2011). The core, however, receives prominent projections from the prelimbic cortex and magnocellular basolateral AMYG as well as small dopaminergic projections from the substantia nigra and functionally plays a role in selectively instigating an approach toward an incentive stimulus associated with the best available reward after Pavlovian cue encoding (Parkinson et al., 2000; Cardinal and Everitt, 2004; Bjorklund and Dunnett, 2007; Basar et al., 2010; Salgado and Kaplitt, 2015; Floresco, 2015).

Similar interspecies microanatomical features and whole-brain axonal projection patterns of subareas in the primate Acb have been qualitatively described (Rigoard et al., 2011; Wedeen et al., 2012; Jbabdi et al., 2013; Balsters et al., 2019), and evolutionarily conserved functions have been suggested for them (Izawa et al., 2003; Calipari et al., 2012; Daniel and Pollmann, 2014) and used as *a priori* knowledge for further studies (Neubert et al., 2015; Heilbronner et al., 2016). However, whether these conserved functions are supported by macroscopic structural connectivity given the disproportionate volumetric changes during primate evolution in the brain regions that project to the Acb (Carlén, 2017; Smaers et al., 2017) and the prefrontal white matter (Schoenemann et al., 2005) has not been established. In addition, researchers found that the striatum, the parent structure of the Acb, presented species differences in regional gene expression (Sousa et al., 2017) and exceptional volumetric changes during primate evolution (Barger et al., 2014). All these seem to support the idea that there may be unique modifications of the structural connectivity of the Acb shell-core architecture, thereby leading to changes in the functions of these Acb subregions.

Quantitative interspecies comparison of structural and functional connectivity has proven to be challenging (Jbabdi et al., 2013; Mars et al., 2014). Compared to traditional comparative anatomy, comparative neuroimaging using magnetic resonance imaging (MRI) allows us to make large-scale comparative analyses and is one of the few techniques that can truly bridge the gap between species by identical non-invasive data acquisition and data handling (Thiebaut de Schotten et al., 2018). Comparative MRI is an effective tool for characterizing multimodal connectivity (Jbabdi et al., 2015; Cloutman and Lambon Ralph, 2012) and has been used to describe species differences in specific fiber pathways (Rilling et al., 2011; Zhang et al., 2013; Hecht et al. 2015; Folloni et al., 2019) and whole-brain connectivity profiles (Mars et al., 2016, 2018a, 2018b; Neubert et al., 2015). Especially for the latter, the connectivity fingerprint (or its variant, e.g., the blueprint) framework has been used to establish a common space in which a brain area’s connectivity profiles characterized in different brains can be aligned and compared between species (Mars et al., 2016, 2018a, 2018b). Such fingerprint-based common space approaches fully exploit the possibilities offered by neuroimaging techniques but have a methodological bottleneck, i.e., how to improve the commonness and representation resolution of the common space to make the approaches more persuasive. For example, there are currently no atlases that allow us to directly extract homologous cortical regions to establish fingerprint-based common spaces for reasonable cross-species comparison.

In this study, we explored the hypothesis that the structural connectivity profile of the Acb shell-core architecture, if characterized by connections with the highly developed PFC regions as well as with the subcortical structures, might differ between humans and macaques because of differences in their evolutionary paths. Compared with mixed comparative approaches (e.g., tracing vs. tractography), as was done in previous studies (Jbabdi et al., 2013; Donahue et al., 2016), we decided that 1) holistic connectivity features-defined brain regions of interest (ROIs) were the best brain nodes for structural and functional connectivity analyses and that 2) using homologous, or at least interspecies connectionally comparable (ICC), regions to characterize the nodes’ connectivity profiles for cross-species comparisons represented a more reasonable approach. Therefore, we first generated the shell-like and core-like regions based on whole-brain tractography in the two species as seeds and delineated pairs of ICC targets based on the similarity between their resting-state functional connectivity (rsFC) profiles. We used these ICC targets to characterize the seeds’ structural connectivity profiles in the two species and compared them using a fingerprint-based common space approach. We also analyzed the possible differences in single structural connectivities.

## Materials and Methods

### Subjects, MRI Data Acquisition, and Overall Processing Flow

We extracted a human MRI dataset (40 subjects; ages: 22-35; 22 males) from the Human Connectome Project (Van Essen et al., 2013) and preprocessed the diffusion (3.0 T MRI scanner; acquired using a multi-shell approach at a 1.25 mm isotropic resolution) and resting-state functional MRI (rsfMRI; 2 mm isotropic resolution; 1,200 time points) data in an earlier study (Xia et al., 2017). In this study, we used these diffusion images to generate the human seeds, i.e., the Acb shell-like and core-like divisions, based on whole-brain voxel-wise tractography (Fig. 1A). Then, we checked the data quality and availability of the rsfMRI data (see Supplemental section 1) and used them to select targets and to delineate the ICC targets (Fig. 1B). Finally, we used the diffusion images to characterize the structural connectivity profiles of the human seeds for cross-species comparisons (Fig. 1C).

**Figure 1.**
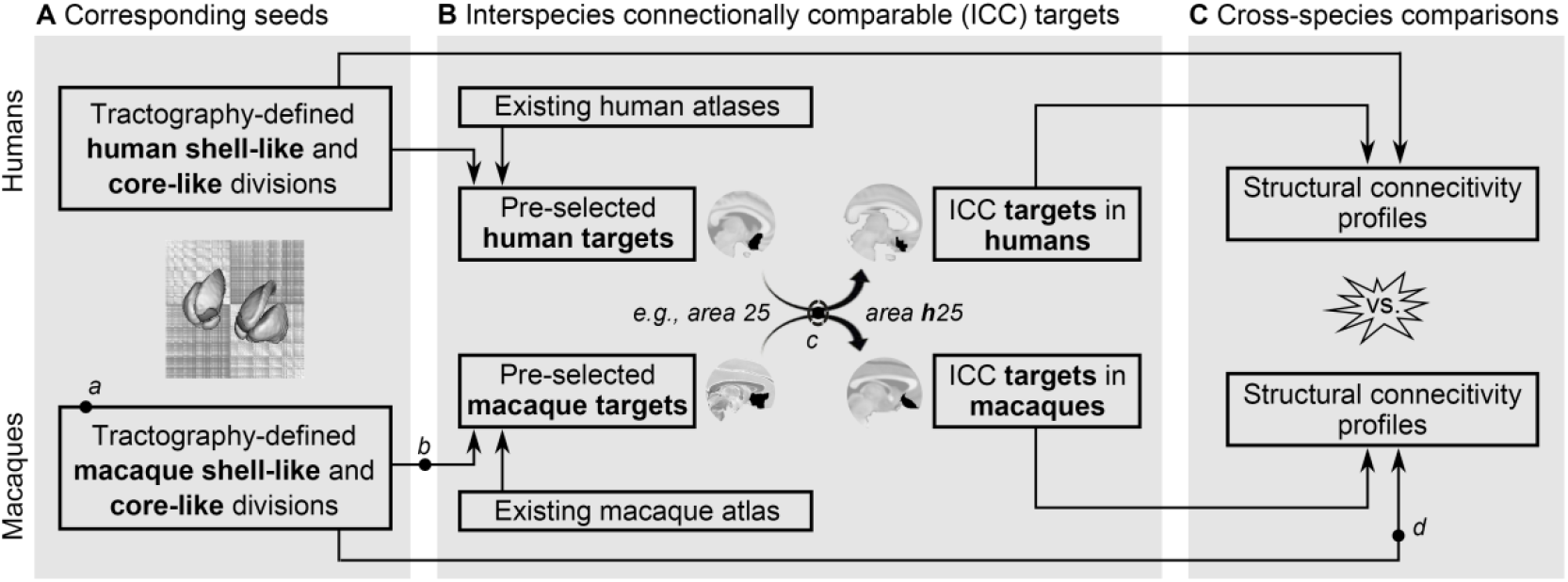
Overall approach of the study. (A) Definition of the seeds. We used whole-brain voxel-wise tractography-based parcellation procedure to generate the human and macaque seeds, i.e., the shell-like and core-like divisions. (B) Definition of the ICC targets in the PFC. Based on the structural and functional connectivity strength with the seeds, we pre-selected many targets from the existing atlases and then redefined their boundaries using a fingerprint-based common space approach to generate the ICC targets for the two species. (C) Cross-species comparisons. We used these ICC targets to characterize the structural connectivity profiles of the seeds in the two primate brains for cross-species comparison. Additional experiments and checks were run on some steps and are marked using ‘*a*, *b*, *c*, and *d*’.

Two rhesus macaque MRI datasets, a high-resolution *ex vivo* macaque MRI dataset (MMDS1; 8 subjects; ages: 4, 4, 5, 6, 8, 12, 15, 23 years; 2 males) and an *in vivo* macaque MRI dataset (MMDS2; 24 subjects; ages: 3.2-4.4 years; body weight: 5.2-6.887 kg; 20 males), from earlier studies (Xia et al., 2019a, 2019b; Wang et al., 2017) were included in this analysis. We had checked the data quality and availability of the low *b*-value diffusion images in MMDS1 (9.4 T animal MRI system; TR/TE = 9800/21.8 ms; voxel sizes = 0.6×0.6×0.6577 mm; 60 diffusion-weighted images, *b* = 1000 s/mm^2^ and 4 non-diffusion-weighted images) to ensure that they could be used to perform the tractographic analysis and used them to define the shell-like and core-like regions based on whole-brain voxel-wise tractography (Xia et al., 2019b). In this study, we extracted these Acb parcels as the macaque seeds (Fig 1A), while using the diffusion images in MMDS2 (3.0 T; voxel sizes = 1.5×1.5×0.65 mm; 63 diffusion-weighted images, *b* = 1000 s/mm^2^ and 1 non-diffusion gradients acquisition) to test the reproducibility of this dichotomous Acb subdivision. Similarity, we checked the data quality and availability of the rsfMRI data in MMDS2 (240 volumes; voxel sizes = 1.803×1.803×1.8 mm; see Supplemental section 1) and used them to define targets and to delineate the ICC targets (Fig. 1B). Finally, we used the diffusion images in MMDS1 to characterize the structural connectivity profiles of the macaque seeds for cross-species comparisons (Fig. 1C). The preprocessing steps (Supplemental section 2) for the macaque MRI data were identical to those used for the human MRI data to mitigate the influence of different preprocessing steps on the cross-species comparison.

### Tractography-defined Seeds of the Shell and Core

Although it is relatively new, tractography-based parcellation is a mature method (Eickhoff et al., 2015, 2018; Fan et al., 2016; see Li et al., 2017 and Supplemental section 3 for details) that has previously been used to generate human and macaque Acb connection units (i.e., a region consisting of voxels with similar structural connectivity features) by parcellating the striatum (Xia et al., 2019a). In this study, we further parcellated these Acb connection units to define the shell and core connection units in the two species. Specifically, we parcellated the human Acb connection unit, rather than the previously used microanatomically delineated Acb region (Baliki et al., 2013; Xia et al., 2017; Zhao et al., 2018), to generate the shell-like and core-like divisions. We had previously parcellated the macaque Acb connection unit into shell-like and core-like divisions based on whole-brain voxel-wise tractography using the high-resolution diffusion images in MMDS1 (Xia et al., 2019b). In this study, we directly extracted these Acb parcels as seeds for the subsequent analysis and repeated this procedure using the low-resolution diffusion images in MMDS2 to validate the reproducibility of the parcellation results (Fig. 1*a*).

### Constructing Fingerprint Frameworks

The fingerprint framework was constructed using a group of targets that met two criteria: 1) have a strong structural connectivity and functional coupling pattern, i.e., rsFC, with either the human or macaque seeds (see detailed criterion in Supplemental section 5) and 2) have substantial evidence-based homologs in this area between the two species. Additionally, the target group should be able to make a unique connectivity characterization of these seeds, so as to distinguish the seeds from each other and from the other neighboring brain regions.

For humans, the prefrontal areas 11, 11m, 13, 14m, 25, 32pl, and 47o extracted from a tractography-defined brain atlas (Neubert et al., 2015) and the AMYG, HIPP, thalamus (THA), midbrain (MidB), pallidum (Pa), caudate nucleus (Ca), and putamen (Pu) extracted from the Harvard-Oxford subcortical atlas (Desikan et al., 2006) were shown to have strong structural and functional connectivities with the shell-like and core-like regions (Xia et al., 2017). For macaques, 8 prefrontal areas, 10m, 11m, 13a, 14m, 14o, 25, 32, and periallocortex, and 7 subcortical structures, the AMYG, HIPP, THA, MidB, Pa, Ca, and Pu, were extracted from a histological atlas (Calabrese et al., 2015; Paxinos et al., 2009). (Note that small subdivisions in this atlas were combined into their parent structure to approximately match the accuracy of the above human atlas.) These prefrontal and subcortical areas were previously shown to have strong structural and functional connectivities with the shell-like and core-like regions (Xia et al., 2019a). In this study, we considered these prefrontal areas and subcortical structures separately as candidates for targets to construct the fingerprint frameworks. In addition, we repeated the above procedure using the histologically-defined macaque shell and core (Paxinos et al., 2009; Calabrese et al., 2015) and validated the reproducibility of the pre-selection of these targets (Fig. 1*b*). Then, we roughly matched these human and macaque target regions based on macroscopic morphological landmarks and added non-matching targets (i.e., corresponding areas selected in only one species) to achieve the first criterion. Next, because some brain areas may have no clear homolog between the two species (Bush and Allman, 2004), we searched earlier studies to determine whether evidence-based non-homologous areas and considerable interspecies differences exist for each pair of targets. If so, the target was eliminated from the group to meet the second criterion.

Because projections from the ventromedial PFC are not limited to the Acb but project broadly in the rostral striatum terminating throughout the medial Ca and Pu (Ragsdale, 1981; Berendse et al., 1992; Haber et al., 1995; Ferry et al., 2000; Haber et al., 2006; Averbeck et al., 2014), we made additional experiments to discover whether both the shell and the core could be distinguished from the rest of the striatum using their structural connectivity profiles characterized by the prefrontal areas as well as by whole-brain voxels (Supplemental section 6). If they did, we could use the prefrontal areas to make unique structural connectivity characterizations of the shell and of the core to perform reasonable cross-species comparisons.

### Delineating ICC Targets in the PFC

The targets identified above for the two species were extracted from different types of brain atlases and could only be roughly matched across the species based on their general location. Specifically, the subcortical targets had relatively clear boundaries. They are undoubtedly pairs of homologs and thus could be used directly to establish the fingerprint-based common space for cross-species comparisons. In contrast, no macroscopic morphological landmarks supported such a clear-cut definition of the homologous target areas in the PFC. So, we did the next best thing: We redefined the boundaries of the pre-selected prefrontal targets based on the similarity of the rsFC profiles to ensure 1) they were connection units and 2) they were ICC brain regions.

Neubert et al. (2015) used 23 evidence-based homologous areas to calculate the rsFC fingerprint of voxels in the human and macaque PFC. Based on the similarity of the rsFC fingerprints between humans and macaques, the authors found pairs of ICC voxels dispersed in this cortical region, i.e., the rsFC profile was used to help relate the two primate brains. In this study, we delineated pairs of ICC targets in the PFC based on the similarity of the rsFC fingerprints calculated using the above 23 homologs (see flow chart in Fig. 2).

**Figure 2.**
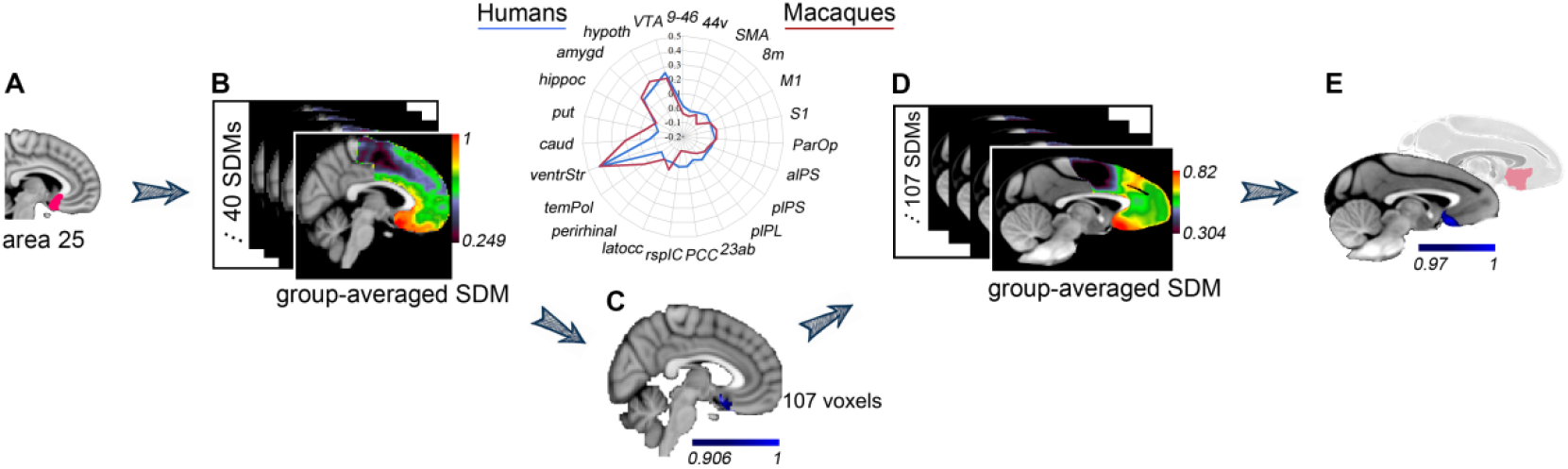
Delineating ICC targets in the PFC. Take area 25 for example: (A) The pre-selected human target of the area 25 extracted from the atlas provided by Neubert et al. (2015). (B) The calculated individual and group-level averaged similarity distribution maps (SDMs). (C) The delineated ICC target in humans (including 107 voxels). (D) A group of SDMs generated in the macaque brain and the group-averaged SDM. (E) The final delineated ICC target in the macaque brain (blue cluster). It is distinctly different from the pre-selected histologically-defined area 25 (red-tinged cluster).

We began the delineation of the ICC targets in the human population, as follows: 1) As suggested by Neubert et al. (2015), we drew 6 mm isotropic ROIs centered on the center-of-gravity of those pre-selected targets (e.g., area 25 shown in Fig. 2A). We brought these ROIs and the 23 homologous areas (6 mm isotropic) from the Montreal Neurological Institute (MNI) space back into native space using the state-of-the-art ANTs’ diffeomorphic transformation model (Avants et al., 2008). A limited number of the voxels that had been misregistered into the cerebrospinal fluid or white matter were moved to their actual locations. Then, we brought them from structural space back into functional space. For each ROI, in each subject’s functional space, 2) we first calculated a series of functional couplings between this ROI and 23 homologous areas to build the rsFC fingerprint, which was considered as the eigen fingerprint. Then, 3) we calculated the rsFC fingerprint for each voxel in the PFC using the 23 homologous areas. Finally, 4) we calculated the cosine similarity between each voxel’s fingerprint and the eigen fingerprint to generate a similarity distribution map (SDM). Voxels in the SDM that are close to 1 reflect a strong possibility of being a member of the connection unit. 5) We transformed all the individual SDMs into MNI space to generate the group-averaged SDM (Fig. 2B), and thresholded it by extracting the top 5% most similar voxels (discrete voxels were removed from the maximal cluster) to obtain the final delineated ICC target in humans (Fig. 2C).

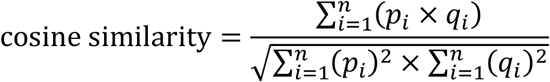

where, *p* and *q* are two vectors representing the functional connectivity fingerprints to be compared; and *n* is the number of arms of the fingerprint.

Given a delineated ICC target in the human brain, we delineated the corresponding ICC target in the macaque brain as follows: 6) We brought the 23 homologous areas (3 mm isotropic; cf. Neubert et al., 2015) from MNI monkey space back into native space (24 macaques in MMDS2) like the first step. 7) For each homologous area, in each subject’s functional space, we first calculated the functional couplings between this area and voxels in the PFC to generate the rsFC map. Then, we transformed these rsFC maps into MNI monkey space to generate the group-averaged rsFC map. The resulting 23 group-averaged rsFC maps were used to extract the fingerprint for each voxel in the PFC in MNI monkey space. 8) For each voxel in the given ICC target in the human brain, we first calculated its fingerprint and considered it as the eigen fingerprint. Then, we calculated the cosine similarities between this eigen fingerprint and the fingerprints of voxels in the macaque PFC to generate a corresponding SDM for the macaque brain. 9) We normalized these SDMs to their respective maximum and then averaged all the SDMs (e.g., 107 SDMs in Fig. 2D). 10) We thresholded the group-averaged SDM by extracting the top 5% most similar voxels (discrete voxels were removed from the maximal cluster) to obtain the final ICC target in the macaque brain (Fig. 2E). Additionally, we adjusted the size of the 23 homologous areas (from 6 mm to 5 mm isotropic in the human brain and from 3 mm to 2 mm isotropic in the macaque brain) to validate the reproducibility of the delineation of the ICC targets (Fig. 1*c*; Supplemental section 8).

### Cross-Species Comparisons

To avoid being influenced by different brain sizes and voxel resolutions from the two species, earlier studies (Mars et al., 2013, 2016; Neubert et al., 2015) used relative connectivity values by normalizing the data to the maximal connectivity value in the brain to build fingerprints for cross-species comparison. However, since the relative connectivity value may be affected by different MRI acquisition parameters, different data processing procedures, and inconsistent normalization factors in different primate brains or even different individual brains within species, we used another connectivity feature, the structural connectivity ratio, to build structural connectivity fingerprints to detect specific structural connectivity differences in the Acb shell-core architecture between the two species.

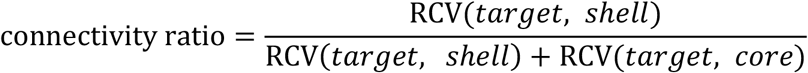

where, *target* is one of the newly delineated ICC targets; RCV(*target, seed*) indicates the relative connectivity value between the *target* and the *seed* (e.g., the shell or core); connectivity ratio thus reflects the degree to which connectivity is biased to the shell or core. It has values in the interval [0, 1], with the high and low values indicating whether the target tended to connect with the shell or the core, respectively.

Similar calculation steps and processing parameters were used to calculate the structural connectivity fingerprints for the two species. More specifically, we brought the two seeds (shell-like and core-like divisions), homologous subcortical targets, and ICC prefrontal targets from MNI space into native space. In the individual diffusion space, we implemented whole-brain tractography for each seed using FSL’s PROBTRACKX2 (50,000 samples; the probability counts were corrected by the length of the pathway; Tomassini et al., 2007) and thresholded these path distribution estimates at *p* > .04% (i.e., 20 out of 50,000 samples; Fan et al., 2016) to limit false positive connections. Then, we calculated the tractographic connectivity probabilities between each target and the two seeds to generate a structural connectivity ratio. Finally, using structural connectivity ratios calculated separately for the prefrontal and subcortical target groups, we calculated fingerprints to characterize the structural connectivity profiles of the Acb shell-core architecture for each individual. In addition, for display purposes, we generated group-averaged structural connectivity fingerprints for the prefrontal and subcortical target groups for the two species.

We used a permutation test to detect whether the structural connectivity profiles of the Acb shell-core architecture are conserved between the two species by evaluating the cosine similarity of their structural connectivity fingerprints. Our null hypothesis assumed that the human and macaque Acb shell-core architecture have conserved structural connectivity profiles. First, we generated the group-averaged tractographic connectivity fingerprints of the human and macaque shell-core architecture and defined their cosine similarity as the observation. Then, we performed the following procedure 1000 times to create the permutation distribution: 1) Two groups of fingerprints were merged and then randomly divided into two groups by keeping their sample sizes static (40 and 8). 2) We generated the group-averaged fingerprints and calculated their cosine similarity. If the null hypothesis was true, the two groups of fingerprints would have identical distributions and the observed cosine similarity would not be rare in the permutation distribution (the test criterion was set at 5%). Thus, we believed there would be no significant interspecies difference between the two groups (humans and macaques) of tractographic connectivity fingerprints. Additionally, a failure to show conservative tractographic connectivity fingerprints could have been caused by one or more significant species differences in a single structural connectivity. Any significant single tractographic connectivity difference between the two species was determined using an independent two-sample *t* test at the 5% significance level.

## Results

### Tractography-Defined Seeds

Both the high-resolution *ex vivo* (Fig. 3A) and low-resolution *in vivo* (Fig. 3B) macaque MRI datasets showed that the Acb can be subdivided into ventromedial and dorsolateral regions based on whole-brain voxel-wise tractography (Xia et al., 2019b). They were named the shell-like and core-like divisions for their corresponding relationships with the well-documented ventromedial shell and dorsolateral core (Fig. 3C). As expected, the parcellation results from the high-resolution MMDS1 group described the data better than those from the low-resolution MMDS2 group (Supplemental section 4), and thus the former were used as the final macaque seeds for subsequent analyses. Similar to the previous parcellation results of the microanatomically defined Acb region (Baliki et al., 2013; Xia et al., 2017; Zhao et al., 2018) and the macaque Acb parcels, the human Acb connection unit could also be subdivided into the ventromedial shell-like and dorsolateral core-like regions based on whole-brain voxel-wise tractography (Fig. 3D). The tractography-defined human and macaque Acb shell-core architecture have similar topological distribution and were considered to be the best brain nodes for subsequent structural connectivity analyses.

**Figure 3.**
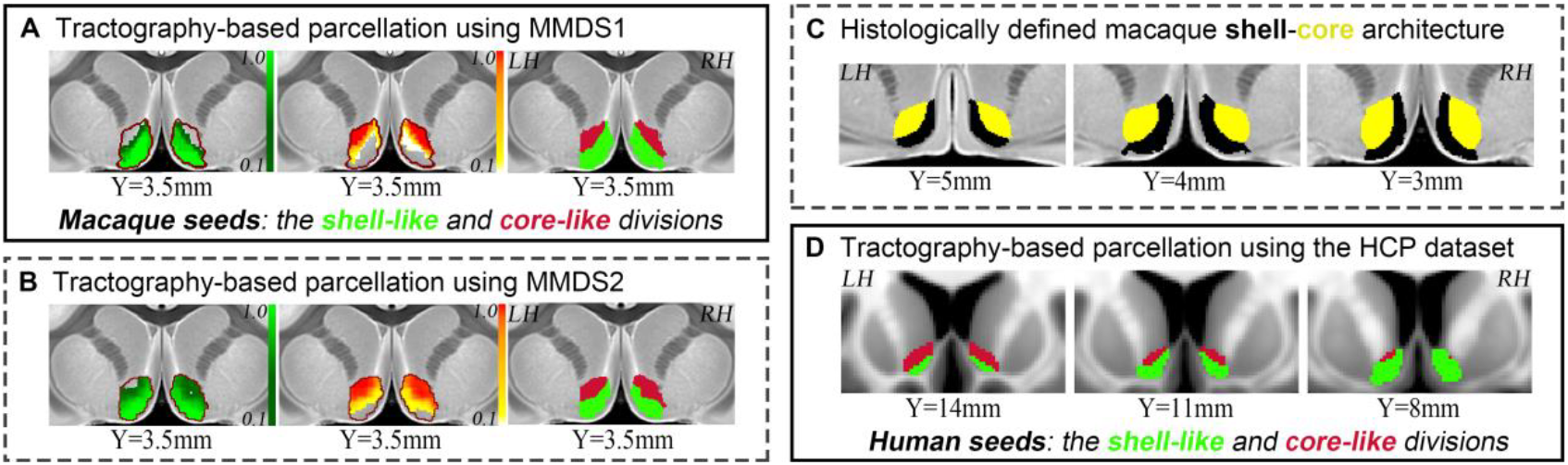
Tractography-defined seeds. (A) The macaque Acb connection unit was parcellated based on whole-brain voxel-wise tractography using the high-resolution *ex vivo* MMDS1 to define the macaque seeds, i.e., the shell-like and core-like divisions. (B) The dichotomous Acb parcels generated by the low-resolution *in vivo* MMDS2. This result ensured the reproducibility of the tractography-defined shell-like and core-like divisions. (C) The histologically defined macaque shell and core provided by Paxinos et al., 2009 and Calabrese et al., 2015. (D) The human Acb connection unit was parcellated based on whole-brain voxel-wise tractography to generate the human seeds, i.e., the shell-like and core-like divisions. Acronyms: left hemisphere, LH; right hemisphere, RH. All the coordinates are shown in MNI (humans) or MNI monkey (macaques) space.

### Defined Fingerprint Framework

The prefrontal areas and subcortical structures enabled us to make unique structural connectivity characterizations of the shell and core (Supplemental section 6). Then, 7 human subcortical structures, the AMYG, HIPP, THA, MidB, Ca, Pu, and Pa, were pre-selected as candidates for the subcortical target group to construct a fingerprint framework. Also, 7 human prefrontal areas, 11, 11m, 13, 14m, 25, 32pl, and 47o, extracted from the tractography-defined atlas (Neubert et al., 2015), were pre-selected as candidates for the prefrontal target group to construct another fingerprint framework. After further investigation, we found that only the most medial part of area 11 (a strip region located in the medial orbital gyrus) had a strong connectivity with the human seeds. We divided this strip region into three and merged these divisions into the neighboring areas 11m, 14m, and 13. Area 47o, located in the posterior part of the lateral orbitofrontal cortex, was eliminated from the target group due to its considerable functional connectivity difference between the two species (Neubert et al., 2015). We confirmed this conclusion in a subsequent delineation of the ICC brain area in area 47o. In the end, we pre-selected 5 prefrontal and 7 subcortical targets in the human brain that could be used to construct fingerprint frameworks. Similarly, using the same analyses for the macaque brain, 7 subcortical structures, the AMYG, HIPP, THA, MidB, Ca, Pu, and Pa, and 8 prefrontal areas, 10m, 11m, 13a, 14m, 14o, 25, 32, and periallocortex, extracted from the histological atlas, were pre-selected separately as candidates for the prefrontal and subcortical target groups to construct the fingerprint frameworks. In addition, we found that the same prefrontal and subcortical targets could be identified when the histologically defined macaque shell and core regions were used as seeds.

Coincidentally, but understandably, the pre-selected 7 human subcortical structures completely overlapped with the pre-selected 7 macaque subcortical structures, i.e., they were identified as being the homologs between the two species and thus could be used directly to construct the final fingerprint framework. On the other hand, the pre-selected human and macaque prefrontal target regions could only be roughly matched based on macroscopic morphological landmarks (Supplemental section 7; Fig. 4). For example, the histologically defined macaque area 32, located in the perigenual anterior cingulate cortex, corresponded to the tractography-defined human area 32pl. Finally, because the number of the pre-selected human prefrontal targets was less than the number of pre-selected macaque prefrontal targets (5 < 8), we began delineation of the ICC targets in the human population.

**Figure 4.**
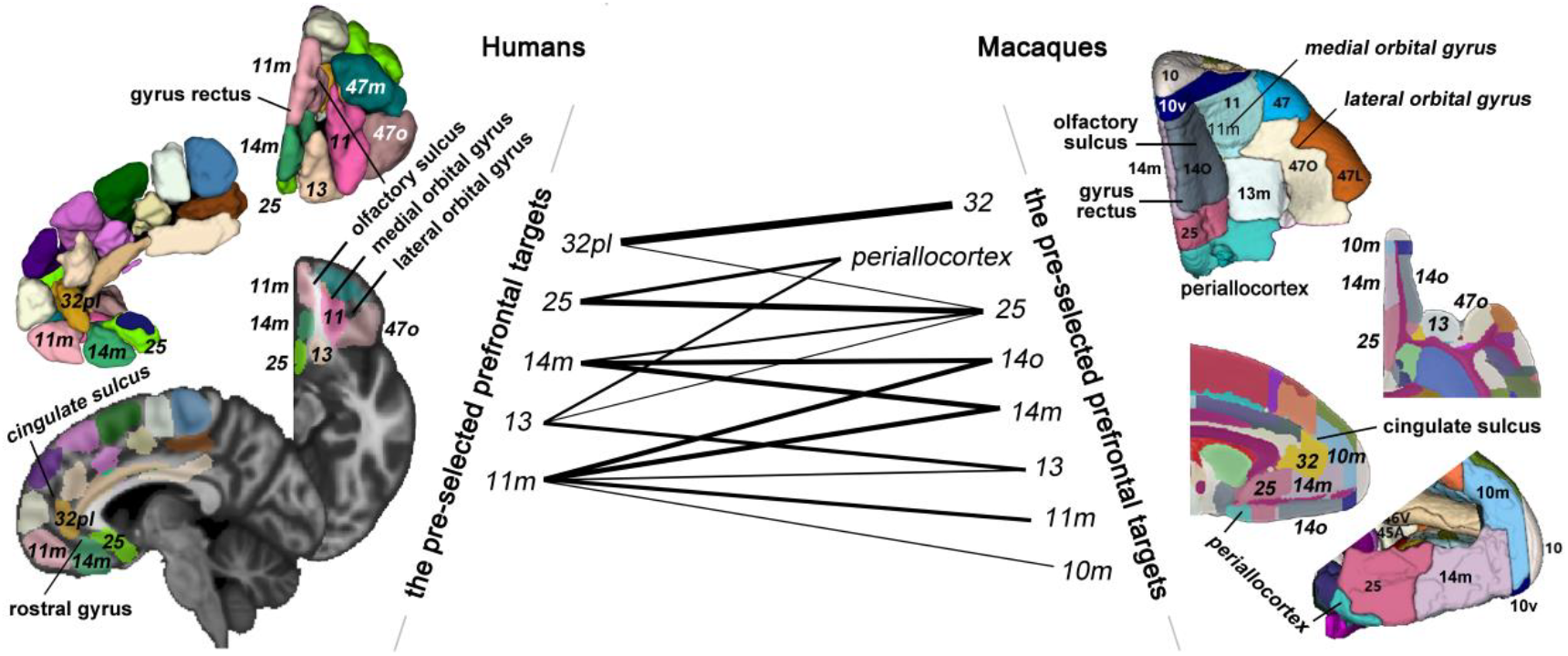
Roughly corresponding relationships of the pre-selected human and macaque prefrontal targets. On the left, we show the pre-selected human prefrontal targets extracted from the tractography-defined atlas (Neubert et al., 2015). On the right, we show the pre-selected macaque prefrontal targets extracted from the histologically defined atlas (Paxinos et al., 2009; Calabrese et al., 2015). We roughly matched these brain areas across the species based on macroscopic morphological landmarks and used thick and thin lines to reflect the strong and slight corresponding relationships, respectively.

### Delineated ICC Targets in the PFC

For each hemisphere, we delineated 5 pairs of ICC targets in the human and macaque PFC. We began this procedure with the 5 human prefrontal targets and thus named them h11m, h13, h14m, h25, and h32 (‘h’ is short for ‘homologous’). Taking **h25** as an example (see the blue cluster in Fig. 5), in each human subject’s functional space, a SDM, summarizing the degree of correspondence between the rsFC patterns of each voxel in the human PFC and the central region of the area 25, was calculated and then transformed into MNI space. We calculated the group-averaged SDM and thresholded it to generate the area h25 in the human brain. Next, for each voxel in the human area h25, we calculated a corresponding SDM in the macaque PFC. Then, the group-averaged SDM was generated and thresholded, with the resulting macaque area h25 being located in histological area 25, with some extensions into the medial periallocortex. Finally, we validated the reproducibility of the delineation of the area h25 (Fig. S5).

**Figure 5.**
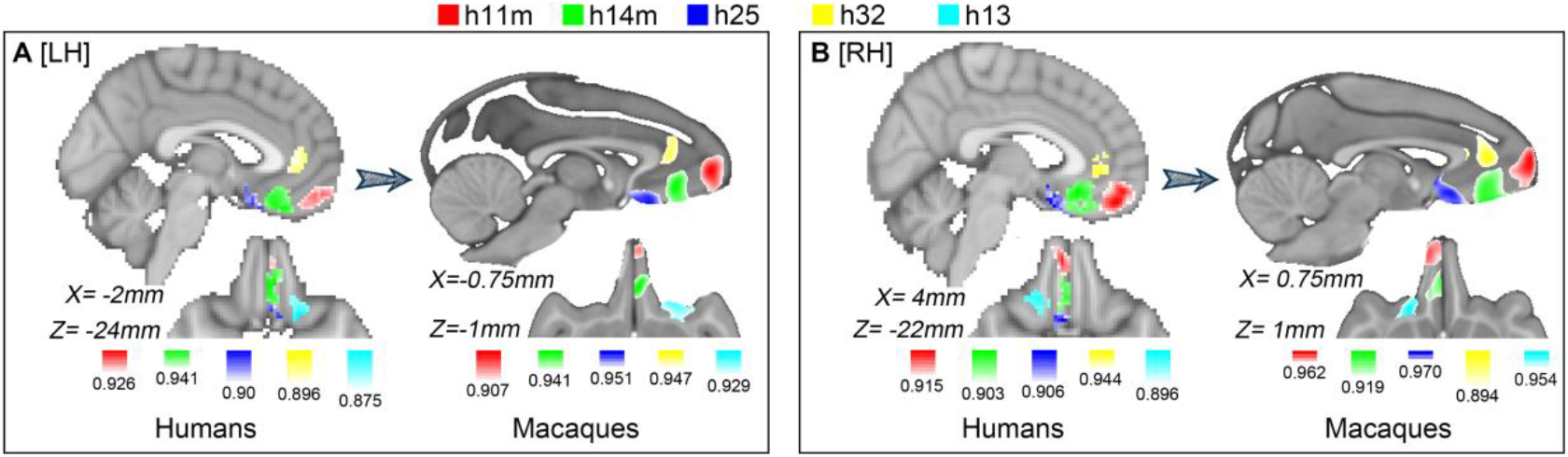
Delineated ICC targets in the PFC. For each hemisphere (A: LH; B: RH), we delineated 5 pairs of ICC targets, h11m, h13, h14m, h25, and h32, which are displayed in red, cyan, green, blue, and yellow, respectively, using the color ranges: [*t*, 1], *t* is the threshold value shown next to the colorbar.

Using this procedure, we delineated other pairs of ICC targets in the two primate brains (Figs. 5, S6, and S7; Supplemental section 8) as follows: the h11m located in the medial frontal pole, the h13 located in the posterior medial orbital gyrus, the h14m located in the middle part of the ventromedial PFC, the h25 located in the posterior part of the ventromedial PFC, and the h32 located in the perigenual anterior cingulate cortex. In addition, we retested the prior description of an interspecies considerable difference in the lateral orbitofrontal cortex by trying to delineate ICC targets using area 47o. No cluster existed in the macaque anterior lateral orbital gyrus after the group-averaged SDM was thresholded.

### Conserved Subcortical Structures-Acb Structural Connectivity

Using the tractography-defined seeds (the shell-like and core-like divisions) and the 7 pre-selected subcortical structures, we built the structural connectivity fingerprint for each individual in the two species and displayed their group-averaged fingerprints in Fig. 6 (top panel). Permutation tests indicated good consistency between the two groups of fingerprints for the two species, as can be seen from the observed cosine similarity, which was less than the calculated test criterion at the 5% significance level in the right tail of the histogram. More specifically, the tractography results indicated that the AMYG and HIPP clearly tended to connect with the shell-like region, a finding that was consistent with earlier imaging results in humans (Baliki et al., 2013; Xia et al., 2017; Zhao et al., 2018) and with tracing findings that the hippocampal projections from the subiculum and CA1 regions were notably restricted to the shell via the fimbria-fornix fiber bundle (Friedman et al., 2002; Poletti and Creswell, 1977). We also found that the caudal basolateral and rostral basal amygdaloid fibers projected throughout the ventral striatum, especially the medial part of the striatum (Friedman et al., 2002; Russchen et al., 1985). Beyond that, we found approximately equivalent structural connectivity ratios (to the shell-like and core-like areas) in the human and macaque brains (see the grouped box charts in Fig. 6). In contrast, the THA, MidB, and Ca tended to connect with the core-like area, a finding which agrees with earlier neuroimaging results in humans (Baliki et al., 2013; Xia et al., 2017; Zhao et al., 2018) and with tracing reports that the shell is distinguished from the core in that it receives the fewest projections from the ventral, anterior, medial, and lateral THA (Gimenez-Amaya et al., 1995) and that the monkeys projections from the Acb (mainly the core) to the substantia nigra in the MidB seem to outnumber those (mainly from the shell) to the ventral tegmental area in the MidB (Haber et al., 1990). We found these regions all had similar structural connectivity ratios between the two primate brains as well.

**Figure 6.**
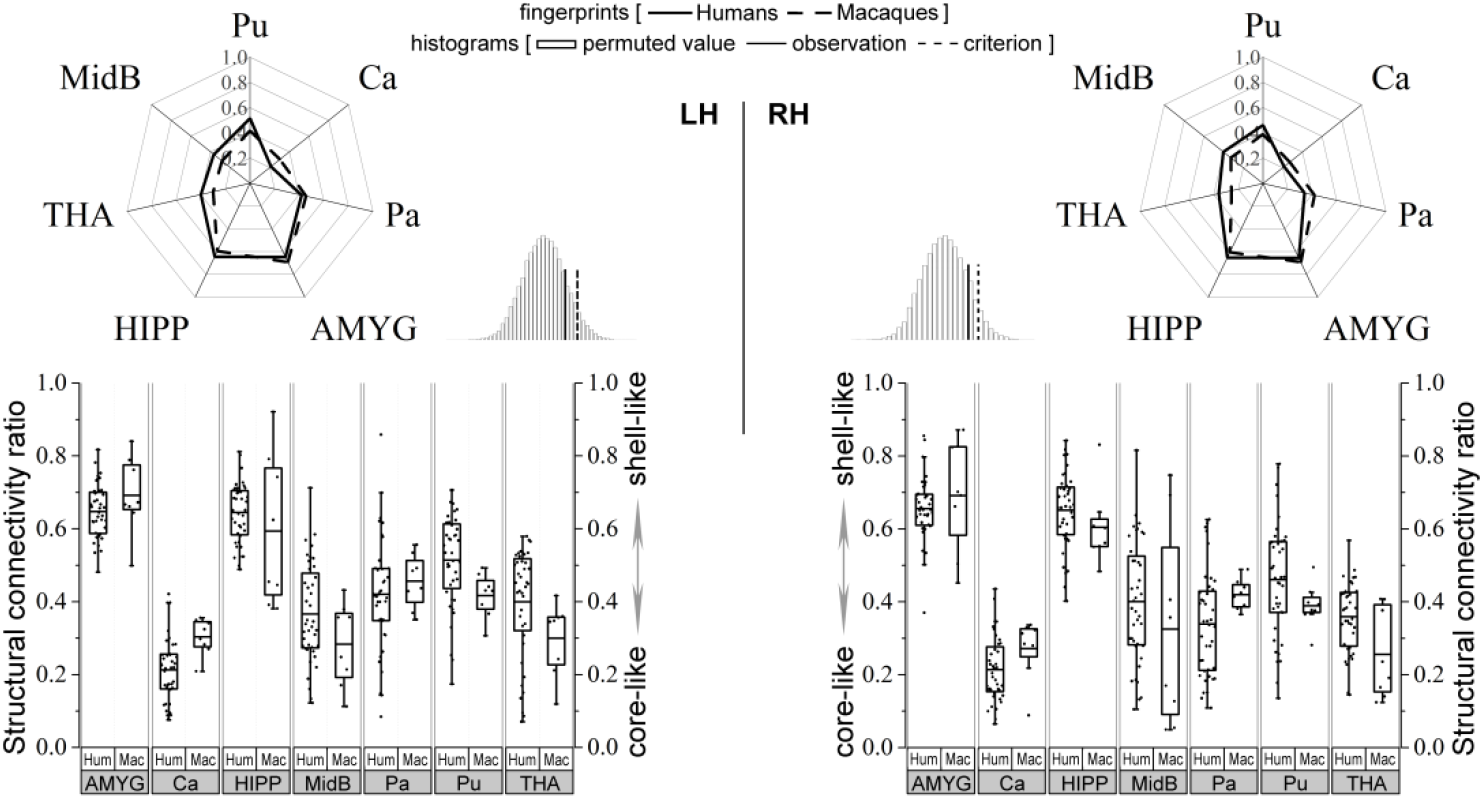
Cross-species comparison of the subcortical structures-Acb structural connectivity. The group-averaged human and macaque structural connectivity fingerprints are shown using radar maps. The convergence between the two groups of fingerprints for the two species was tested using a permutation test, and the results are shown in the histogram. The single structural connectivity ratio was calculated for each subcortical target in the two species. An independent two-sample *t* test was used to determine whether the two population means were equal, and the result is shown in the grouped box chart. The results indicate that the observed cosine similarity between the two group-averaged fingerprints of the two species was less than the calculated test criterion in the right tail of the histogram, suggesting a similar distribution of the two groups of fingerprints. In addition, no statistically significant single structural connectivity ratio difference was found between the two species. Acronyms: putamen, Pu; caudate nucleus, Ca; pallidum, Pa; amygdala, AMYG; hippocampus, HIPP; thalamus, THA; midbrain, MidB; Humans, Hum; Macaques, Mac.

### Altered PFC-Acb Structural Connectivity

Using these seeds and 5 delineated ICC prefrontal targets, we built additional pairs of structural connectivity fingerprints for each individual in the human and macaque groups and displayed the two group-averaged fingerprints in Fig. 7 (top panel). Unlike the above subcortical result, we found a rare cosine similarity in the permutation distribution, as can be seen in the right tail of the histogram. The observed cosine similarity was higher than the calculated test criterion at the 5% significance level. Therefore, we rejected the null hypothesis and concluded that the prefrontal areas of the two primate species evolved dissimilar gross structural connectivity trends to the shell-core architecture. This result seems to align with evidences for heterochronic developmental shifts in the human PFC and prefrontal white matter during primate evolution (Carlén, 2017; Smaers et al., 2017; Schoenemann et al., 2005; Somel et al., 2011). We concluded that one or more prefrontal targets had noticeably different structural connectivity trends in the two species and that this could be further checked by performing single structural connectivity analyses.

**Figure 7.**
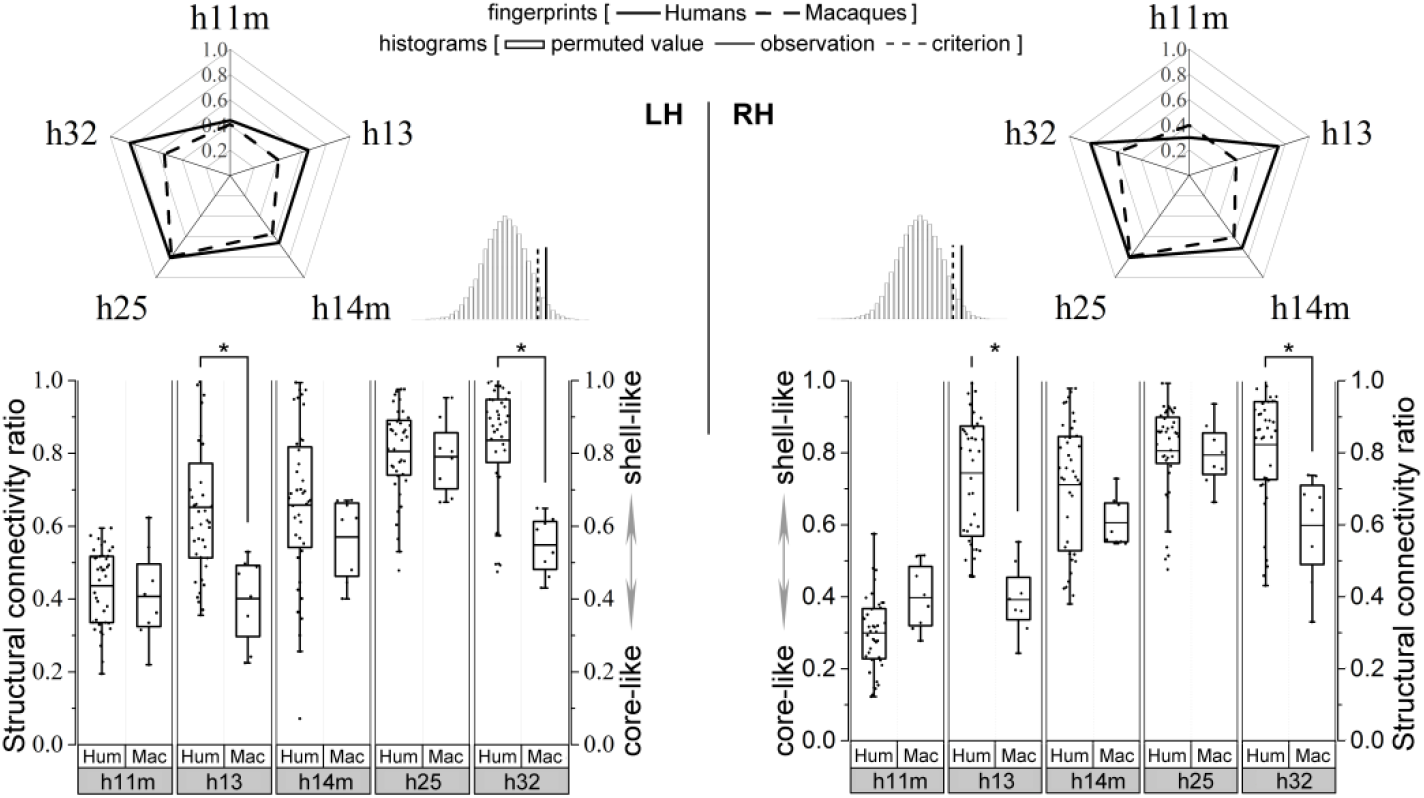
Cross-species comparison of PFC-Acb structural connectivity. Please refer to Figure 6 for a detailed legend. The results indicate that the two groups of fingerprints could not be accepted as convergent fingerprints because the observed cosine similarity was higher than the calculated criterion in the right tail of the histogram. The gross structural connectivity ratio (i.e., fingerprint) differences between the two species were localized to h13 (LH: humans: 0.61 ± 0.20 > macaques: 0.40 ± 0.12; *p* = 5.7e-03. RH: humans: 0.74 ± 0.17 > macaques: 0.39 ± 0.10; *p* = 1.65e-06) and h32 (LH: humans: 0.82 ± 0.16 > macaques: 0.55 ± 0.08; *p* = 3.62e-05. RH: humans: 0.79 ± 0.16 > macaques: 0.60 ± 0.15; *p* = 3.1e-03). Both of these showed significantly different structural connectivity ratios (* *p* < .05).

For each delineated prefrontal target, we calculated the structural connectivity ratio between this region and the two seeds in each of the two species (Fig. 7). More specifically, area h25 had a stronger structural connectivity to the shell-like area than to the core-like (about 0.8 of the structural connectivity ratio) in both species, which was consistent with previous findings that the rodent infralimbic area (rodent homolog of the primate posterior ventromedial PFC) was the main origin of cortical projections to the Acb and was notably restricted to the medial shell (van Kuyck et al., 2007). Moreover, we found no interspecies differences in the connectivity ratio of the h25 (and the other two areas in the ventromedial PFC, the h14m and h11m) to the two Acb parcels between the two species, suggesting that during primate evolution connectional strengthening or weakening of these prefrontal pathways to the shell-core architecture occurred approximately proportionately in the two types of primate brains. In addition, previous tracing results indicated that the ventromedial PFC and lateral orbitofrontal cortex mainly targeted the shell and core, respectively (Ferry et al., 2000; Haber et al., 1995). We obtained a similar result from neuroimaging and also found that the prominent tractographic connectivity between the shell-like area and h25 extended into the neighboring h13 in humans but not in macaques. Similar to the findings of a previous study (Baliki et al., 2013), we found no preferred structural connectivity of h32 with the core-like areas in either of the two species, which seems to contradict the tracing result (Basar et al., 2010; Salgado and Kaplitt, 2015). However, we did find that the relative connectivity values between the two Acb subregions and h32 and their connectivity ratio all appeared significantly stronger in macaques than in humans (for detailed comparison results using relative connectivity values, see Supplemental section 10).

## Discussion

We generated spatially corresponding shell-like and core-like divisions as seeds and delineated ICC targets in human and macaque brains to build structural connectivity fingerprints for viable cross-species comparisons. We revealed that the prefrontal but not the subcortical target group had dissimilar structural connectivity with the shell-core architecture between the two species. We localized this overall difference to specific prefrontal targets in single structural connectivity analyses and analyzed their possible influence on functions.

Microstructural and molecular features (e.g., cyto- and myelo-architecture) may determine the local processing capabilities of a brain region, whereas connectivity governs the nature and flow of information (Hilgetag and Grant, 2000; Passingham et al., 2002; Johansen-Berg et al., 2004). Many studies have thus suggested that function may depend more on connectivity than on microstructural features (Cloutman and Lambon Ralph, 2012; Knosche and Tittgemeyer, 2011). In addition, although tractography does not exactly reflect the measures that traditional tract tracing does (Borra and Luppino, 2018), significant advances in our understanding of brain anatomy have already been made by using this technology because of its ability to show the brain connectivity (Grandjean et al., 2017; see Supplemental section 9 for a simple comparison between the two technologies). Based on these, in this study, we used tractographic connectivity characteristics to characterize and compare the shell-core architecture between humans and macaques to supplement previous comparative findings, such as similarities in the cellular and molecular composition and distribution in the shell and core between the species (Volkow and Morales, 2015; Graybiel and Ragsdale, 1978; Meredith et al., 1996).

Fingerprint-based common space approaches fully exploit the possibilities offered by neuroimaging techniques and have been used in recent comparative MRI studies (Mars et al., 2016, 2018a; Neubert et al., 2015). In this study, some measures were used to enhance the reasonability of the common space approaches. First, a series of connection units, including tractography-defined seeds and rsFC-defined ICC targets, were generated to support the subsequent connectivity analyses. Second, to understand whether the connections between homologous brain regions have changed, it helps to be able to overlay homologous brain regions of the species of interest and then compare their connectivity matrices (Thiebaut de Schotten et al., 2018). In view of this, we identified the homologous relationships of the pre-selected subcortical targets in the two species and redefined the boundaries of pre-selected prefrontal targets to make sure they were ICC regions. In addition, we validated the reproducibility of the definition of the seeds, pre-selected the targets, delineated the ICC prefrontal targets, and characterized the structural connectivity of the shell-core architecture (Supplemental section 11). The verification results indicated that there was no serious snowballing effect resulting from the chain analysis used in this study. In any case, all these measures enabled us to establish a viable comparative framework while holding many other irrelevant factors constant so that we could obtain credible comparison results.

Many evolutionarily conserved structural and functional features of the subcortical structures, including the Acb, have been suggested (Izawa et al., 2003; Calipari et al., 2012; Daniel and Pollmann, 2014; Balsters et al., 2019) and used as *a priori* knowledge (Heilbronner et al., 2016; Neubert et al., 2015). But researchers have also pointed out that many of their features had changed because of their different evolutionary paths. For instance, the AMYG and HIPP show disproportionate volumetric growth and the striatum (which may influence the Acb) shows shrinkage in the human brain compared with what would be predicted based on primate scaling trends. Based on these findings, some researchers have suggested that many of the functions of these homologous structures may not necessarily be conserved (Barger et al., 2014; Stephan, 1983; Stephan et al., 1987). In this study, by showing that the fingerprints had a strong cosine similarity, we revealed that the structural connectivity profiles of the shell-core architecture characterized by the subcortical substructures appears to be practically unaffected by their disproportionate volumetric changes. This interspecies similar white matter layout is consistent with many single conserved projections that have been suggested by tracing studies (Friedman et al., 2002; Leong et al., 2016). These include the amygdaloid and hippocampal projections to the Acb and the spiral-like organization of the striato-mesencephalic-striatal projections. Our result suggests that any subtle structural connectivity variations in these single connectivities between the two species have not shifted the overall structural connectivity profiles of the shell-core architecture.

Heterochronic genetic and developmental shifts of the human PFC regions (Somel et al., 2011; Smaers et al., 2017) have also brought macro-evolutionary scale volumetric changes to these regions, e.g., the disproportionate volumetric growth of the prefrontal gray and white matter areas (Schoenemann et al., 2005; Smaers et al., 2017; Carlén, 2017) and their structural connectivity differences with the shell-core architecture revealed in this study. Also, these structural connectivity differences may be accompanied by some functional modifications, which we inferred and summarized as follows: 1) Area h13 has been indicated as related to identifying reward outcomes in humans (Klein-Flügge et al., 2013) and to encoding the context-invariant value of goods in monkeys (Padoa-Schioppa and Assad, 2008). These distinct functions may be respectively supported by their distinctive preferred structural connectivity with the human shell-like and the macaque core-like regions. 2) Area h32 has been implicated as involved with working memory for action-outcome-based sequencing (Coutureau and Killcross, 2003). In addition, the projection from this area to the striatal (mainly in the core) patch pathway has been identified as causally required for decision-making in conflict situations (Friedman et al., 2015). Significantly stronger structural connectivity in macaques than in humans may suggest a greater ability of macaques to adjust or switch their reward-related actions in conflict situations. In contrast, similar interspecies’ single structural connectivity of brain areas in the ventromedial PFC to the shell-core architecture may suggest some conserved functions, such as the reward and decision-making process related those neural projections (Floresco, 2015; Basar et al., 2010; Salgado and Kaplitt, 2015).

In conclusion, we improved the fingerprint-based common space approaches by providing a reasonable comparative framework. Using this approach, we revealed conserved structural connectivity profiles of the Acb shell-core architecture with subcortical structures but dissimilar structural connectivity profiles with prefrontal regions. These results are in keeping with the well-accepted conclusion that the human PFC has undergone considerable expansion and become more developed than that of other primates. However, all these conserved and changed structural connectivities only reflect general neuroanatomical evolutionary trends. Whether or not any task-specific-related function of the Acb subregions has been affected will, of course, be partially dictated by which connections are involved and how the signals are integrated and interact via these connections.

## Supporting information

Supplemental section 1

## Funding

This work was supported by the Natural Science Foundation of China (Grant Nos. 91432302, 31620103905, 81501179, and 61976150), the Science Frontier Program of the Chinese Academy of Sciences (Grant No. QYZDJ-SSW-SMC019), National Key R&D Program of China (Grant No. 2017YFA0105203), Beijing Municipal Science & Technology Commission (Grant Nos. Z161100000216152, Z161100000216139, and Z181100001518004), Beijing Advanced Discipline Fund, and the Natural Science Foundation of Shanxi Province of China (Grant No. 201801D121135). Human MRI data were provided by the Human Connectome Project, WU-Minn Consortium (Principal Investigators: David Van Essen and Kamil Ugurbil; 1U54MH091657) funded by the 16 NIH Institutes and Centers that support the NIH Blueprint for Neuroscience Research.

## Notes

We thank Shan Yu and George Paxinos for their insightful discussions and useful suggestions. We also thank Rhoda E. and Edmund F. Perozzi for editing assistance.

## Conflict of Interest

None declared. Results are available for download from Gitlab at https://gitlab.com/xlxia/cross-species-comparison-of-the-Acb.

## Notes

### Competing Interest Statement

The authors have declared no competing interest.

