## Supplemental section 1 for "Mapping Connectional Differences between Humans and Macaques in the Nucleus Accumbens Shell-Core Architecture"

Xia et al.

### Supplemental Experimental Procedures and Results

#### (1) Quality Checking of the rsfMRI data

Although the HCP rsfMRI data had a longer scan time and higher voxel resolution, which significantly improved the images' SNRs and thus the temporal SNR (tSNR, dividing the mean of a time series by its standard deviation). We still assessed the quality of the data to make sure they can delineate interspecies connectionally comparable (ICC) targets in the PFC based on the similarity of the voxel-wise resting-state functional connectivity (rsFC) profile. Of course, the same assessing process was also performed on the macaque rsfMRI data in MMDS2. We found undesired interactions with pulsations in the orbitofrontal cortex and the cerebrospinal fluid (CSF) in the ventricles in the two species, which resulted in a reduced temporal stability of the MR signal around these regions. Despite this issue, the majority of voxels showed a higher tSNR than the estimated criterion (Murphy et al., 2007), i.e., humans: about 160 > 42; and macaques: about 68 > 40 based on parameters such as the voxel volume (humans: 8 $mm^3$; macaques: 5.83 $mm^3$) at 3 Tesla (Fig. S1).

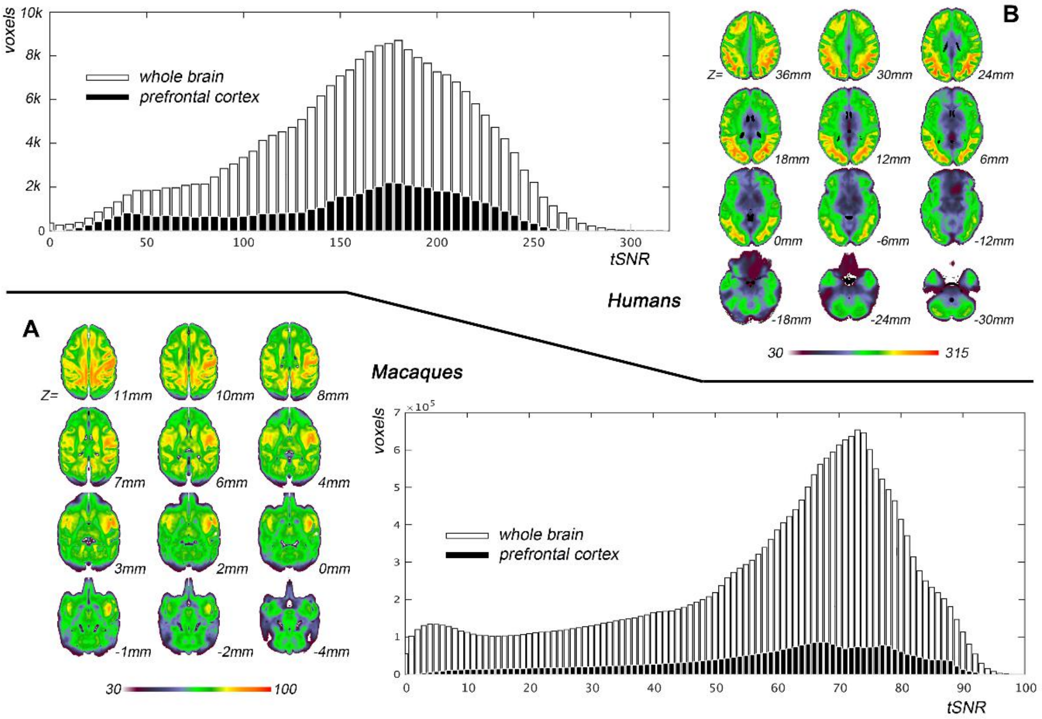

**Figure S1.** Average tSNR across all acquisitions for the two species. In the histograms of the average tSNR scores, the lighter shaded histogram shows the tSNR distribution of all the voxels in an approximate brain mask; the dark shaded histogram shows the tSNR distribution of all the voxels in the prefrontal cortex. Multi-slice images show the spatial distribution of the average tSNR across the brain.

#### (2) MRI Data Preprocessing

The macaque structural images were preprocessed as follows: 1) A correction for the distortion induced by inhomogeneity was made. 2) Because the extraction of the brain can be a problem for the *in vivo* macaque brain data, a rough brain extraction was made to register the data to the INIA19 brain template (Rohlfing et al., 2012). The resulting affine matrix together with their provided tissue (gray matter, GM, white matter, WM, and CSF) probability maps were used to generate relatively accurate subject-native tissue probability maps using FSL's FAST (Zhang et al., 2001) and corresponding binary images (GM, WM, and CSF were thresholded at .4, .99, and .99, respectively). Some poorly recognizable regions in the thalamus (THA) and midbrain (MidB) were corrected manually. 3) The subject-native brains were resampled and rotated to align with the anterior commissure-posterior commissure of the brain template in Montreal Neurological Institute (MNI) monkey space (Frey et al., 2011) with FSL's 6 degrees of freedom (DOF) FLIRT. 4) The subject-native brains were then registered to the brain template using ANTs' Symmetric Normalization transformation model (Avants et al., 2008) to generate the transformation files. 5) All subject-native GM masks were transformed to MNI monkey space for the calculation of the group-averaged GM mask (thresholded at > .5).

These macaque diffusion images (MMDS1 and MMDS2), same with the human data, were preprocessed as follows: 1) A correction was performed to remove the distortion caused by eddy currents and simple head motions. 2) The brain mask was co-registered into subject-native diffusion space using the 6 DOF FLIRT boundary based registration algorithm (BBR, Greve and Fischl, 2009). 3) BEDPOSTX (2 fibers per voxel; Jbabdi et al., 2012) was used to build up the distributions of the diffusion parameters at each voxel for the subsequent probabilistic tractography. The macaque rsfMRI images (MMDS2) were preprocessed as follows: 1) The first 10 volumes were discarded. 2) The brain mask was co-registered into subject-native functional space to remove non-brain tissue. 3) Each volume underwent slice-timing correction. 4) Motion correction of the time-series was performed using MCFLIRT (Jenkinson et al., 2002). 5) Spatial smoothing was performed using FSL's SUSAN with a Gaussian 3 mm FWHM kernel. 6) Detrending was done to remove linear and quadratic trends. 7) Independent component analysis based denoising was performed using MELODIC. 8) A band-pass filter was used to separate the data at slow-4 (.027-.073 Hz; Zuo et al., 2010) to reduce low-frequency drift and high-frequency noise. Note that because of the difference in the voxel resolution (both in absolute number and relative to the size of the brain) of the scans of the two species' brains, we chose a Gaussian 3 mm and a 6 mm FWHM kernel to smooth the macaque and human rsfMRI data, respectively.

##### (3) Tractography-Based Parcellation

The steps of the Tractography-based parcellation procedure, in short, included:

1) The seed mask was brought from standard MNI or MNI monkey space back into each subject's structural space using the symmetric diffeomorphic image registration (Avants et al., 2008). After minor manual modifications of the voxels mis-registered into the WM and CSF, this mask was then brought back into individual diffusion space. In each subject's diffusion space, 2) whole-brain probabilistic tractography was implemented for each voxel in the mask using PROBTRACKX2 (Behrens et al., 2007) by sampling 50,000 streamlines to estimate the connectivity probability. Note that the probability counts were corrected by the length of the pathway to compensate for the distance-dependent bias (Tomassini et al., 2007). 3) These path distribution estimates were thresholded at $p > .04\%$ (i.e., 20 out of 50,000 samples) as was done in earlier studies (Fan et al., 2016; Xia et al., 2017; Li et al., 2017) to limit false positive

connections. 4) All the connectivity probability maps were formed into a connectivity matrix. 5) A cross-correlation matrix between the connectivity profiles of all the voxels in the seed mask was calculated (Johansen-Berg et al., 2004) and was then 6) fed into normalized-cut spectral clustering to subdivide these voxels into multiple subgroups based on the similarity of the whole-brain voxel-wise connectivity profiles (Baldassano et al., 2015). 7) The voxels in each subgroup were mapped back onto the brain to generate the corresponding subregion. 8) All individual parcellation results were transformed into standard MNI or MNI monkey space. In the standard space, for each solution, 9) the most consistent labeling scheme across subjects was adopted to resolve the cluster label mismatch issue caused by the random labeling of the clustering algorithms. Then, 10) groups of locationally corresponding subregions were extracted to generate the probability maps for the subregions. The maximum probability map of the seed was calculated by assigning each voxel of the reference space to the area in which it was most likely to be located (Wang et al., 2012).

##### (4) Definition of the Seeds

We used the above tractography-based parcellation to define the human and macaque seeds. Unlike previous studies (Baliki et al., 2013; Xia et al., 2017; Zhao et al., 2018), we parcellated the Acb connection unit (a tractographic connectivity-defined Acb-like region provided by Xia et al. 2019), instead of the microanatomically defined Acb region, to make sure the final shell-like and core-like divisions were connection units. In addition, we found that the parcellation results from the high-resolution MMDS1 group described the data better than those from the low-resolution MMDS2 group (Fig. S2A). We also found that the tractography-defined and histologically defined Acb shell-core architectures have similar topological distribution as well as some differences (Fig. S2C).

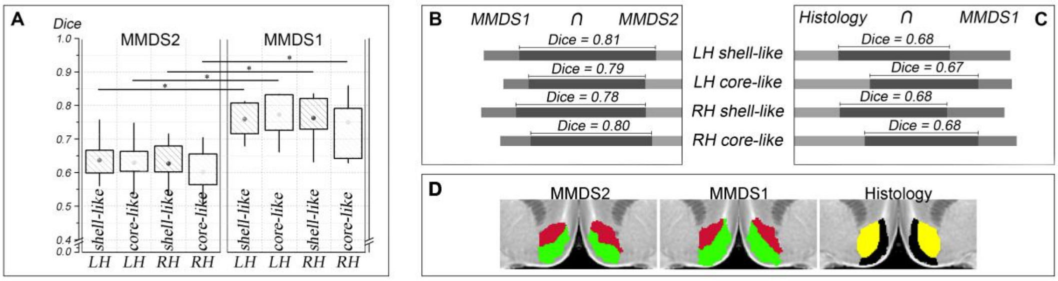


**Figure S2.** Comparisons between Acb parcellation results. (A) The inter-individual consistency (Dice) of the parcellation results from MMDS1 were significant higher than those from MMDS2 (two-sample *t* test: left hemisphere, LH shell-like region: .77 ± .06 > .63 ± .06; core-like region: .76 ± .05 > .64 ± .06; right hemisphere, RH shell-like region: .75 ± .08 > .60 ± .08; core-like region: .76 ± .07 > .63 ± .08; * $p$ < .001). (B) The Dice overlaps between the Acb subregions generated from the two macaque datasets. (C) The Dice overlaps between the tractography-based Acb subregions and corresponding histological results. (D) Different Acb parcellation results.

#### (5) Definition of the Targets

The dichotomous Acb subregions were brought back into each subject's diffusion and functional spaces. For each Acb subregion, in each subject's diffusion space, the whole-brain tractographic connectivity probability map was generated and then thresholded to reduce false positive connections. All individual maps and their binary images were transformed into MNI or MNI monkey space. These individual maps were used to generate a group-averaged tractographic connectivity probability map and their binary images were used to generate a probability fiber tract map. As in previous studies (Fan et al., 2016; Xia et al., 2017), to further reduce false positive connections and the effects of individual differences, the probability fiber tract map was thresholded at $p$ > 50% to generate the common fiber tract map. This step is, in effect, analogous to a one-sample *t*-test to determine the voxels that have significant tractographic connectivity with the given Acb subregion. Finally, the averaged tractographic connectivity probability map was masked by the common fiber tract map to generate the final averaged tractographic anatomical connectivity probability map of this Acb subregion. Referring to earlier literature (Cauda et al., 2011; Xia et al., 2017), we chose brain area which had strong tractographic connectivity with most of the Acb subregions as a member of the target group. We used the averaged structural connectivity probability map to define strong tractographic connectivity according to either of the following two criteria:

1) The number of activation voxels included in the area surpassed a fixed fraction, > 2%, of the total voxels of that area.

2) The number of activation voxels included in the area surpassed a fixed fraction, > 2%, of the total number of activation voxels.

For each subregion, in each subject's functional space, the Pearson correlation coefficients between the mean time series of the given subregion and the time series of each voxel in the GM mask was calculated to generate the rsFC map. Note that the mean time series of these Acb subregions were calculated using fMRI data without smoothing. The rsFC map was converted to $z$-values using Fisher's $z$-transformation and transformed into MNI monkey space. All the normalized $z$-valued rsFC maps were fed into a random effects one-sample $t$-test to determine the regions that had significant correlations with the given subregion. A statistical threshold of $p < .05$ was set to achieve a corrected cluster-wise statistical significance of $p < .05$. The cluster size was estimated on the basis of the GM mask and the group-averaged Gaussian filter width. Then, a minimum statistic test for conjunction (Nichols et al., 2005) was performed among these Acb subregions so that the surviving voxels had significant rsFC with all the Acb subregions. The extended threshold for the cluster size of the conjunction was set at 50. Then, to avoid too many targets causing the risk of overfitting the fingerprint (Mars et al., 2016), we defined a tighter criterion to choose chose brain area which had strong rsFC with most of the Acb subregions as a member of the target group. We used the significant rsFC map to define strong rsFC according to either of the following two criteria:

1) The number of the active voxels included in the area surpassed a fixed fraction of total voxels of that area at > 5%.

2) The number of the active voxels included in the area surpassed a fixed fraction of total activation voxels at > 5%.

**(6) Usability Analysis of the Target Groups**

Projections from the ventral and medial prefrontal cortex  are not limited to the core. Rather, they project broadly in the rostral striatum terminating throughout the medial caudate nucleus and putamen (Pu) across species. In view of this, we preformed experiments and analyses to decide whether the prefrontal areas can make a unique characterization of the structural connectivity profiles of the shell and the core to distinguish them from each other and from the rest of the striatum. We had parcellated the striatum based on whole-brain voxel-wise tractography to identify the Acb-like region and its neighboring regions, e.g., the ventral Pu (Xia et al., 2019; see Fig. S3A). Then, we used a group of prefrontal areas (10, 13, 14, 25, and 32) to characterize the region-wise structural connectivity fingerprint of the Acb-like and ventral Pu. After comparison, we found that the two regions have clearly distinct structural connectivity profiles (Fig. S3B). Furthermore, our earlier study (Xia et al., 2017) indicated that the Acb shell-like and core-like divisions can be distinguished by their unique structural connectivity with a group of prefrontal areas (11, 11m, 13, 14m, 23ab, 24, 25, 32d, 32pl, 47o, 47m, and FPm). In short, we think that the region-wise connectivity profile represented by prefrontal areas can be used to distinguish the shell and core from each other and from the rest of the striatum.

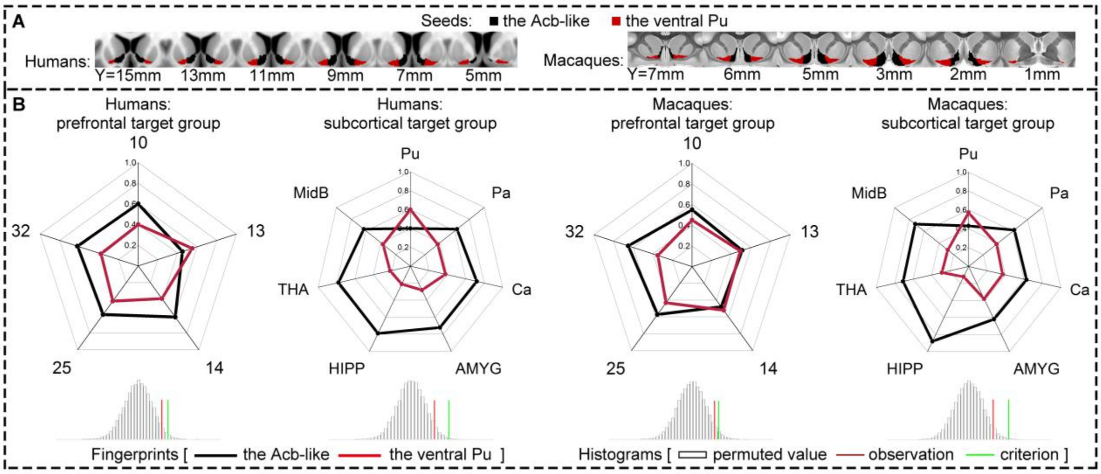


**Figure S3.** The unique structural connectivity profile of the Acb-like and ventral Pu regions. (A) The human and macaque Acb-like and ventral Pu regions are shown in a multi-slice presentation. (B) The group-averaged structural connectivity fingerprints for the Acb-like and ventral Pu regions are shown using the radar maps. The permutation tests results are shown using the histograms.

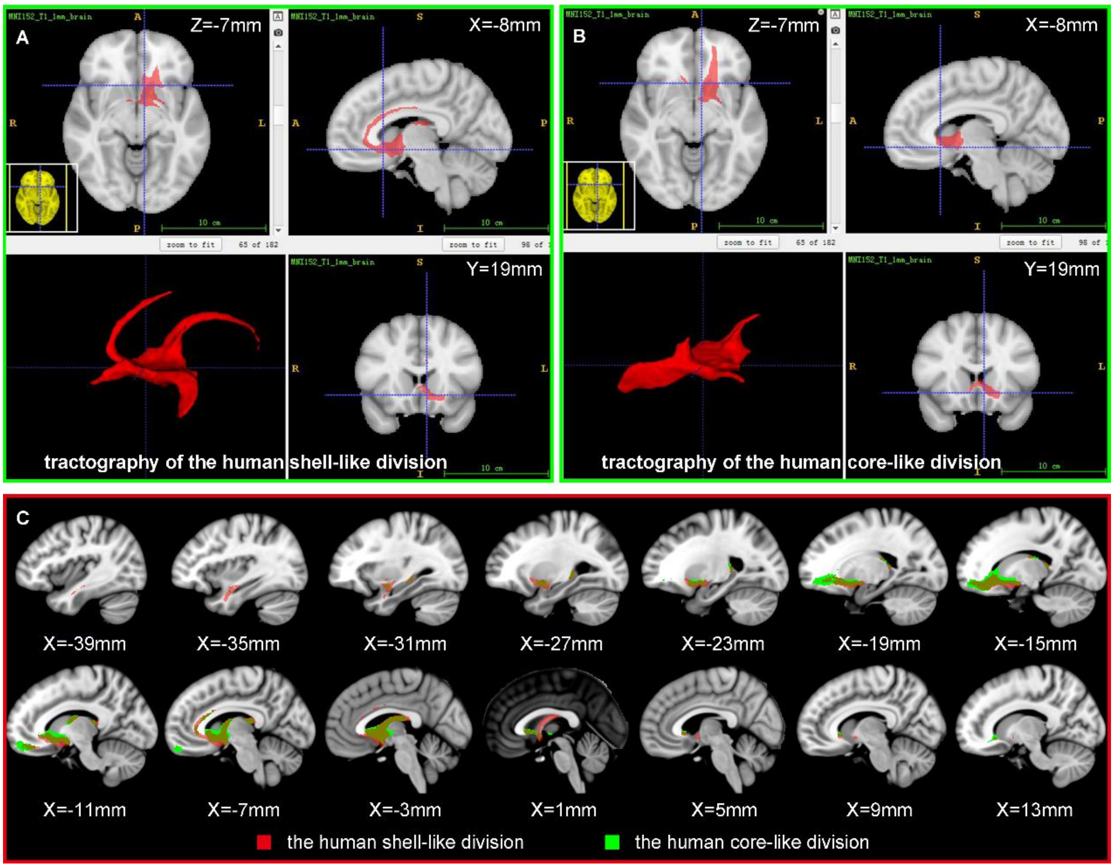

**Figure S4.** The whole-brain probabilistic tractography of the human shell-like and core-like divisions. For display purposes, the group-averaged probabilistic tractography of the shell- and core-like regions were thresholded at 70% and are portrayed on MNI152 template in 3D and multi-slice presentations.

Previous studies have indicated that the human shell- and core-like divisions can be identified based on whole-brain voxel-wise structural and functional connectivity, and co-activity patterns (Baliki et al., 2013; Xia et al., 2017; Zhao et al., 2018). We further characterized their whole-brain voxel-wise structural connectivity (see Fig. S4) and region-wise structural connectivity fingerprints using a group of prefrontal areas (Xia et al., 2017). We found a significant difference between the fingerprints of the shell-like and core-like divisions and thus suggested that the prefrontal areas could be used to make a unique connectivity characterization of the shell and the core.

#### (7) Rough Regional Corresponding Relationship Between the Two Species

Different brain atlases might be defined by different features and methods. In such situations, the labeled areas in different atlases may only be roughly matched based on macroscopic morphological landmarks rather than others features across the species. In this study, based on the strong structural and functional connectivity with the Acb subregions, we identified many prefrontal areas in humans and macaques and roughly matched them by macroscopic morphological landmarks, as follows: The connection between the macaque area 10m and the Acb subregions was primarily focused on its ventral part. This region, located in the anterior part of the gyrus rectus, corresponded to human area 11m. Macaque area 11m, located in the anterior medial orbital gyrus (MOG), corresponded to human merged area 11m (including area 11m and anterior medial area 11). Macaque area 13a, located in the posterior MOG (most of the area was covered by area 13 and the orbital periallocortex), corresponded to human merged area 13 (including area 13 and posterior medial area 11). Macaque area 14m, located in the rostral gyrus and medial part of the gyrus rectus, corresponded to human areas 14m and 11m. Macaque area 14o, located in the medial part of the gyrus rectus with an extension into the MOG, corresponded to human merged area 14m (including area 14m and the middle part of medial area 11) and merged area 11m. Macaque area 25, located in the rostral gyrus and the posterior part of the gyrus rectus, corresponded well to human area 25 with an extension into merged area 14m. Macaque orbital periallocortex, located in the posterior part of the gyrus rectus, corresponded to human area 25 with an extension into merged 13. Macaque area 32, located in the perigenual anterior cingulate cortex, corresponded to human area 32pl.

##### (8) Delineated ICC Targets in the PFC

We took area h25 as an example to validate the reproducibility of the ICC brain areas. We used 23 evidence-based homologs (3 mm isotropic macaque ROIs and 6 mm isotropic human ROIs) to generate the averaged similarity distribution map, SDMs, of the area h25 in the two primate brains (Fig. S5A). Then, we adjusted the volume size of the 23 homologs (2 mm isotropic macaque ROIs and 5 mm isotropic human ROIs) to generate the group-averaged SDMs of the area h25 in the two primate brains (Fig. S5B). We indicated that the group-averaged SDMs of the human and macaque area h25 were slightly affected by the two different sizes of the 23 homologs. The final 10 pairs of ICC PFC targets calculated by 3 mm isotropic macaque ROIs and 6 mm isotropic human ROIs are shown in Figs. S6 and S7 and are described in Table S1.

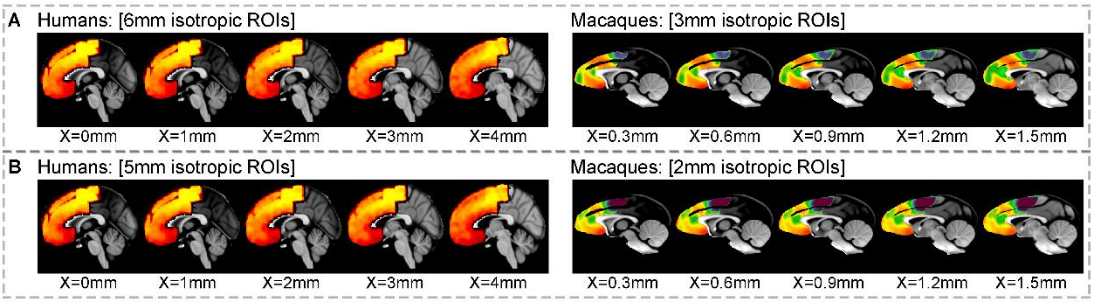


**Figure S5.** Reproducibility of the delineation of the area h25. (A) The SDMs of area h25 delineated using 6 mm isotropic human ROIs and 3 mm isotropic macaque ROIs were similar to corresponding (B) SDMs of area h25 delineated using 5 mm isotropic human ROIs and 2 mm isotropic macaque ROIs.

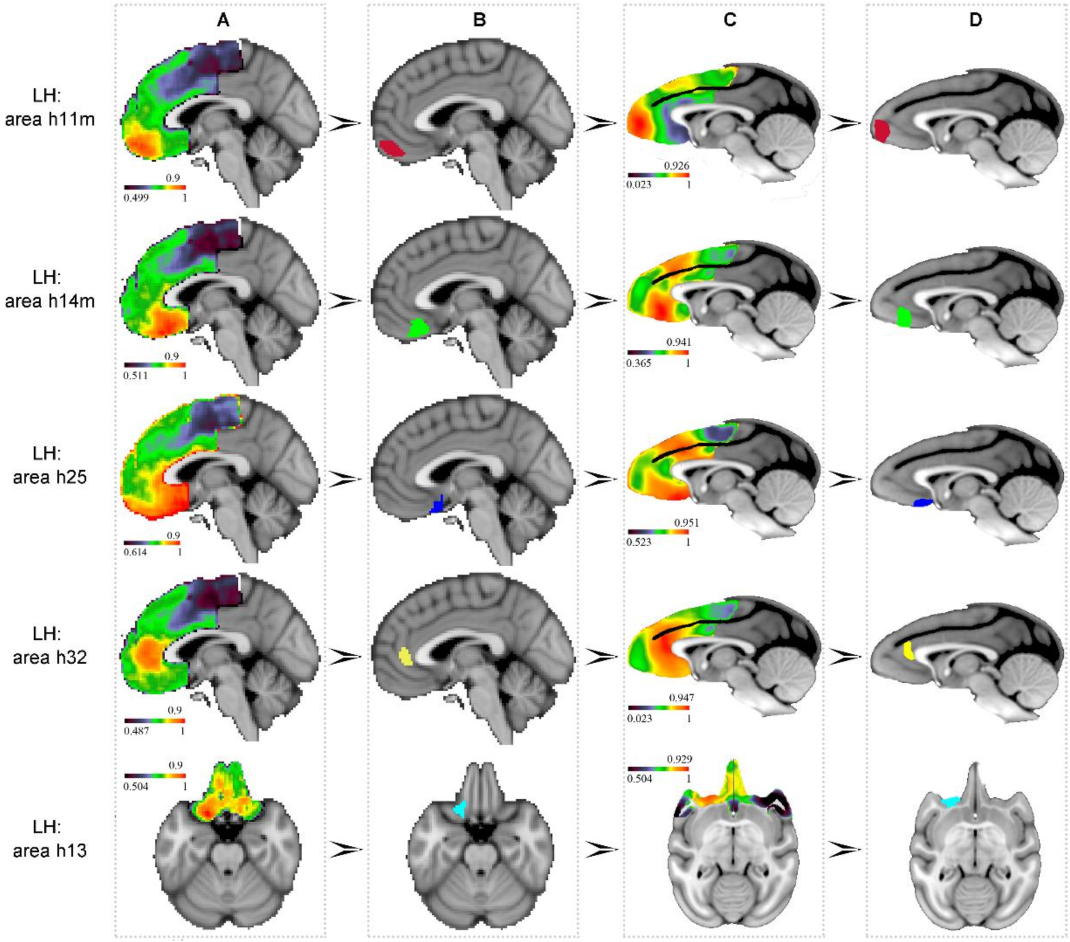


**Figure S6.** ICC targets in the left PFC. The four-column images from left to right indicate (A) the group-averaged SDM in the human brain -> (B) the final ICC targets in the human brain -> (C) the group-averaged SDM in the macaque brain -> (D) the final ICC targets in the macaque brain. Each ICC target was generated by the threshold *t* shown next to the colorbar.

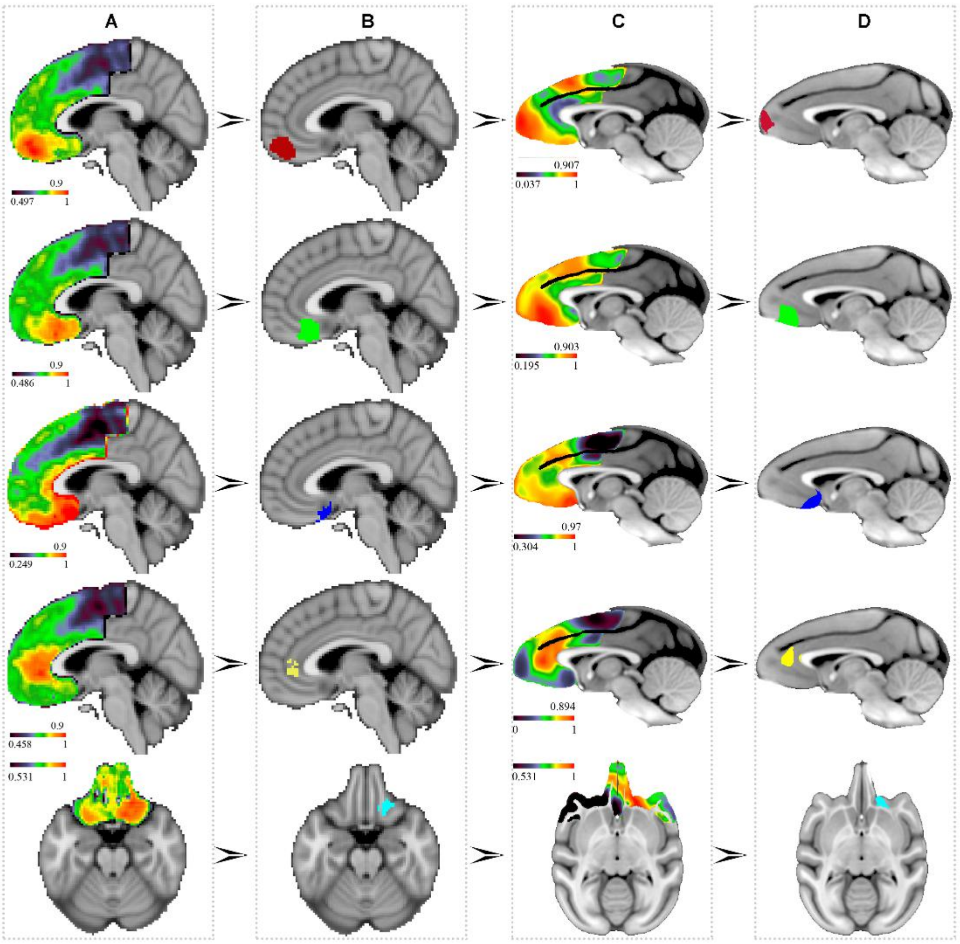


**Figure S7.** ICC targets in the right PFC.

**Table S1.** Descriptive information for the ICC targets.

| Region | Humans | | Macaques | |
|---|---|---|---|---|
| | Volume | Center of gravity | Volume | Center of gravity |
| LH: h11m | 1528 $mm^3$ | [-4.54, 52.58, -15.02] | 76 $mm^3$ | [-1.87, 23.91, 0.78] |
| LH: h14m | 2000 $mm^3$ | [-3.58, 31.89, -18.60] | 95 $mm^3$ | [-1.64, 16.12, -1.81] |
| LH: h25 | 984 $mm^3$ | [-2.96, 12.46, -18.81] | 25 $mm^3$ | [-0.92, 9.33, -3.50] |
| LH: h32 | 2384 $mm^3$ | [-6.21, 44.03, 5.09] | 67 $mm^3$ | [-1.95, 14.44, 6.99] |
| LH: h13 | 1424 $mm^3$ | [-18.00, 22.24, -19.46] | 106 $mm^3$ | [-9.42, 11.29, 0.50] |
| RH: h11m | 2192 $mm^3$ | [4.28, 51.77, -16.64] | 111 $mm^3$ | [2.24, 24.53, 2.20] |
| RH: h14m | 2056 $mm^3$ | [3.46, 30.05, -17.21] | 140 $mm^3$ | [1.84, 16.62, -1.65] |
| RH: h25 | 888 $mm^3$ | [5.17, 14.70, -18.68] | 44 $mm^3$ | [0.99, 8.07, -2.60] |
| RH: h32 | 1736 $mm^3$ | [8.53, 42.80, 2.52] | 98 $mm^3$ | [1.19, 15.50, 6.08] |
| RH: h13 | 1592 $mm^3$ | [18.56, 26.58, -18.21] | 63 $mm^3$ | [8.26, 11.62, 0.22] |

#### (9) Comparisons Between Tract Tracing and Tractography

We made a simple comparison between tract tracing and tractography (Fig. S9). We generated the voxel-wise tractographic connectivity probability maps of the shell-like and core-like regions (Fig. S8) and used them to calculate the regional connectivities between the Acb subregions and targets (Fig. S9B). We compared the results with the previous tracing results provided by Ferry et al. (2000) (Fig. S9A). The comparison results showed that although tractography does not exactly reflect the measures that tract tracing do, the connectivity sites and connectivity trend are similar between the two measuring technologies.

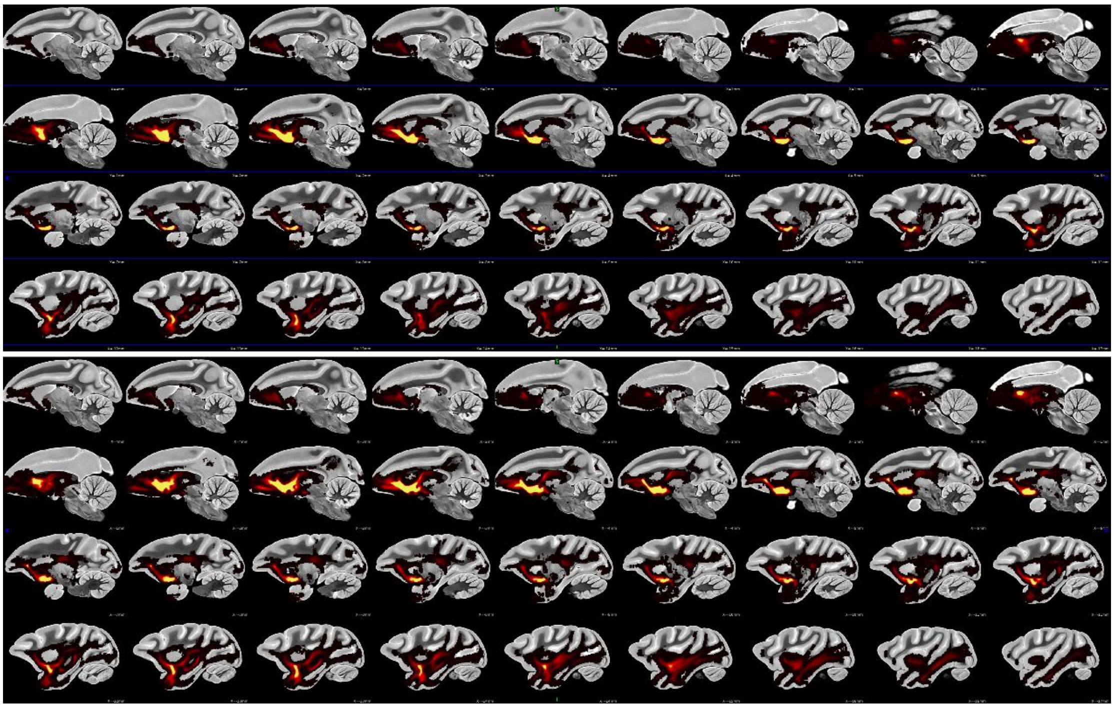

**Figure S8.** The whole-brain voxel-wise tractographic connectivity probability maps of the macaque Acb shell-like (top panel) and core-like (low panel) regions.

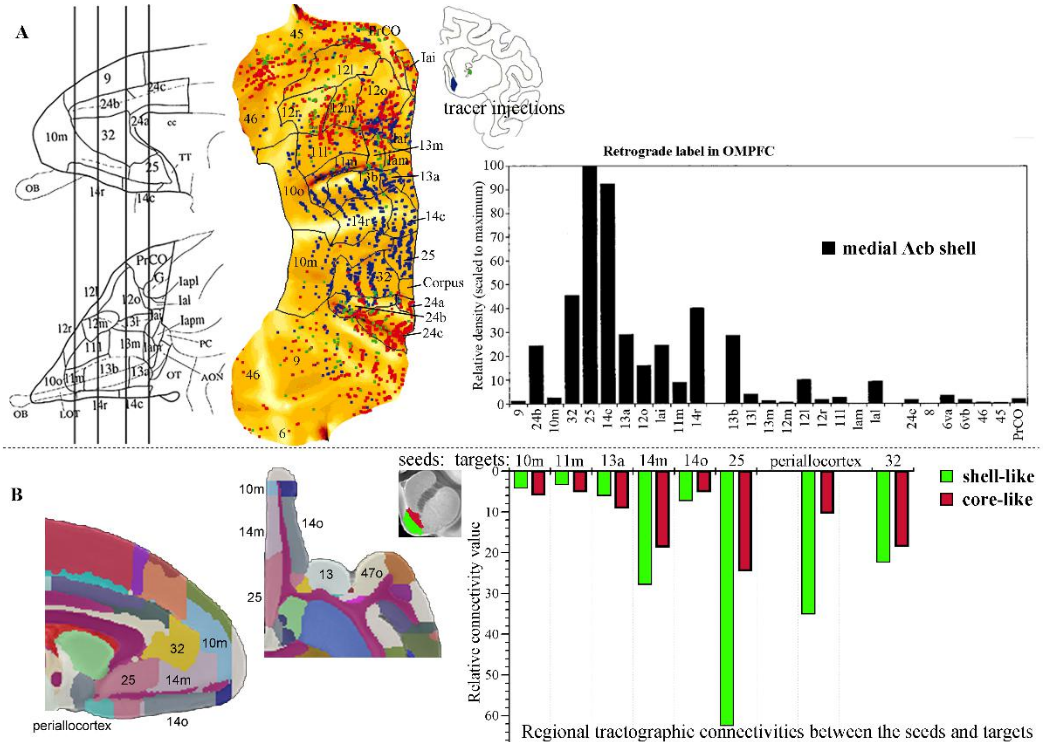


**Figure S9.** Comparison of tractography and tract tracing measures of connectivity strength in rhesus macaque. (A) The relative density of retrogradely labeled cells in the macacca fascicularis orbital and medial PFC (atlas: Szabo and Cowan, 1984) after retrograde tracer injection in the medial shell was provided by Ferry et al. (2000). Bar graph indicates the relative density of retrogradely labeled cells normalized to the greatest density. Their result showed that the medial shell receives projections mainly from gyrus rectus (area 25, 14c, 14r, and 11m), perigenual anterior cingulate cortex (area 32 and 24b), and medial MOG (area 13a, 13b, and lai). (B) In this study, the relative tractographic connectivity values were calculated between the Acb subregions and the identified prefrontal targets extracted from the rhesus macaque atlas (Paxinos et al., 2009). Our result showed that the shell mainly connect with the gyrus rectus (area 25, periallocortex, 14m, 14o, 11m, and 10m), perigenual anterior cingulate cortex (area 32), and medial MOG (area 13a) as well.

#### (10) Cross-Species Comparison Using the Regional Relative Connectivity

We calculated the relative connectivity between each Acb subregion and ICC target by normalizing the data to the whole-brain averaged tractographic connectivity in the two species to investigate species structural connectivity strength differences.

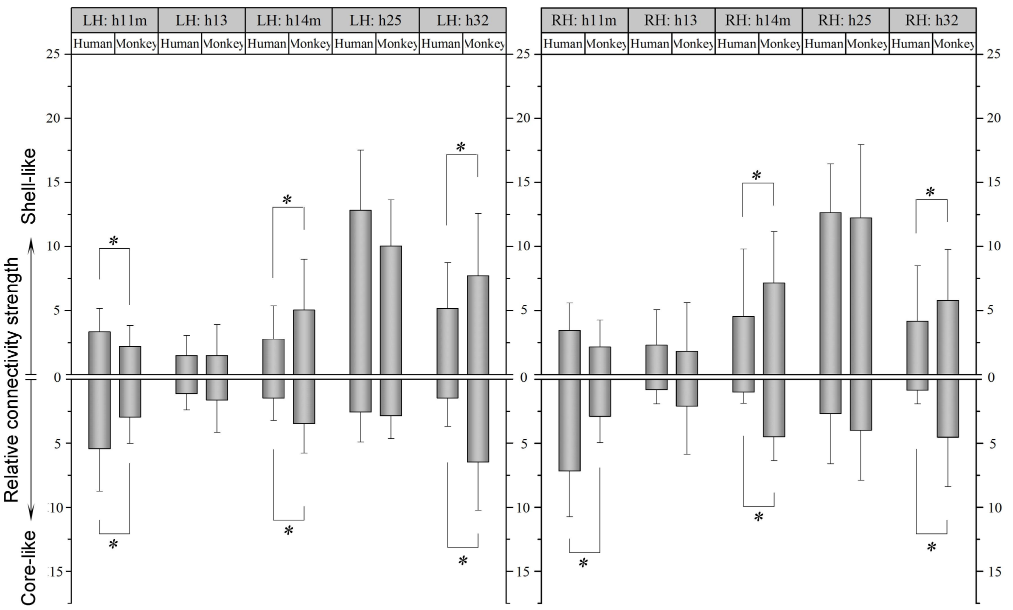


**Figure S10.** Cross-species comparisons of single structural connectivity. The two species had different regional tractographic connectivity to the shell-like (top panels) and core-like region (bottom panels), which were localized to the h11m, h14m, and h32 (independent two-sample *t* tests were used in this procedure; * p < .05).

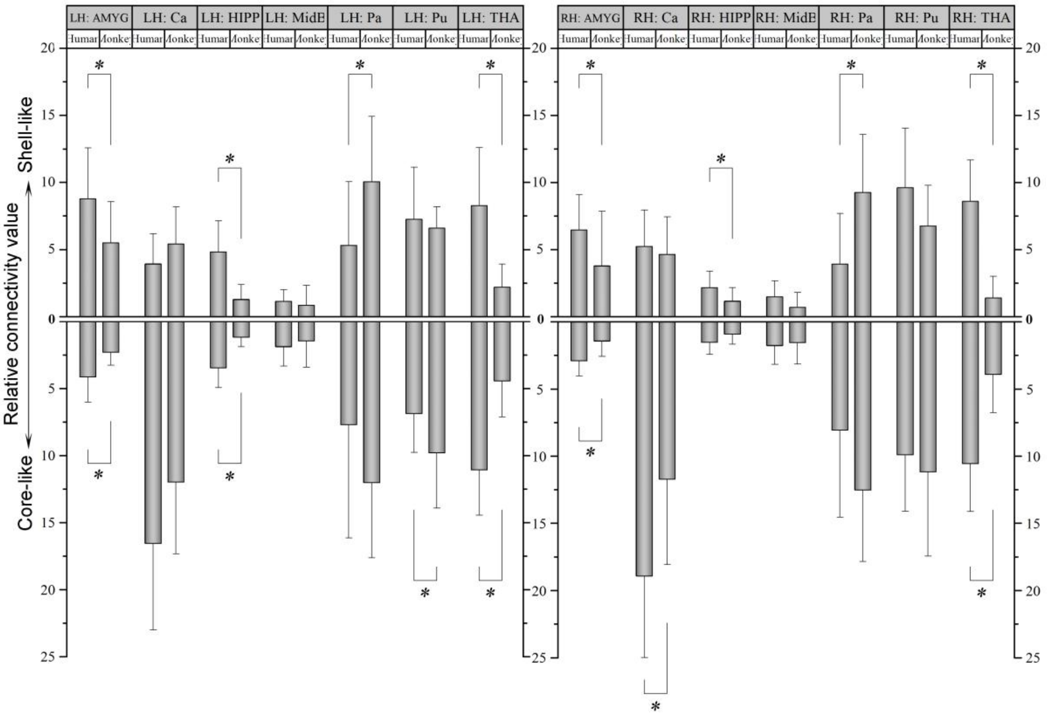

**Figure S11.** Cross-species comparisons of single structural connectivity. The two species had different regional tractographic connectivity to the shell-like (top panels) and core-like region (bottom panels), which were localized to many subcortical structures, e.g., the AMYG, HIPP, Pa, and THA (* $p < .05$).

#### (11) Reproducibility Tests

We ran additional experiments and checks on main steps, including the definition of the seeds (Fig. 1*a*), the pre-selection of the targets (Fig. 1*b*), and the delineation of the ICC PFC targets (Fig. 1*c*), to validate the reproducibility of the results to indicate that a serious snowballing problem did not exist. Besides the above, we also validated the reproducibility of the characterization of the structural connectivity of the shell-core architecture using the histologically defined seeds in the macaque brain (Fig. 1*d*). For each target, we calculated and grouped the individual structural connectivity ratios using the tractography- and histologically-defined shell-core architecture (see the box charts in Fig. S10B). Two-sample *t* tests at the 5% significance level were used to test the significance of the differences between corresponding ratio groups. However, we did not find any target that showed a (tractographic) structural connectivity ratio that was significantly different from the tractography- and histologically-defined seeds.

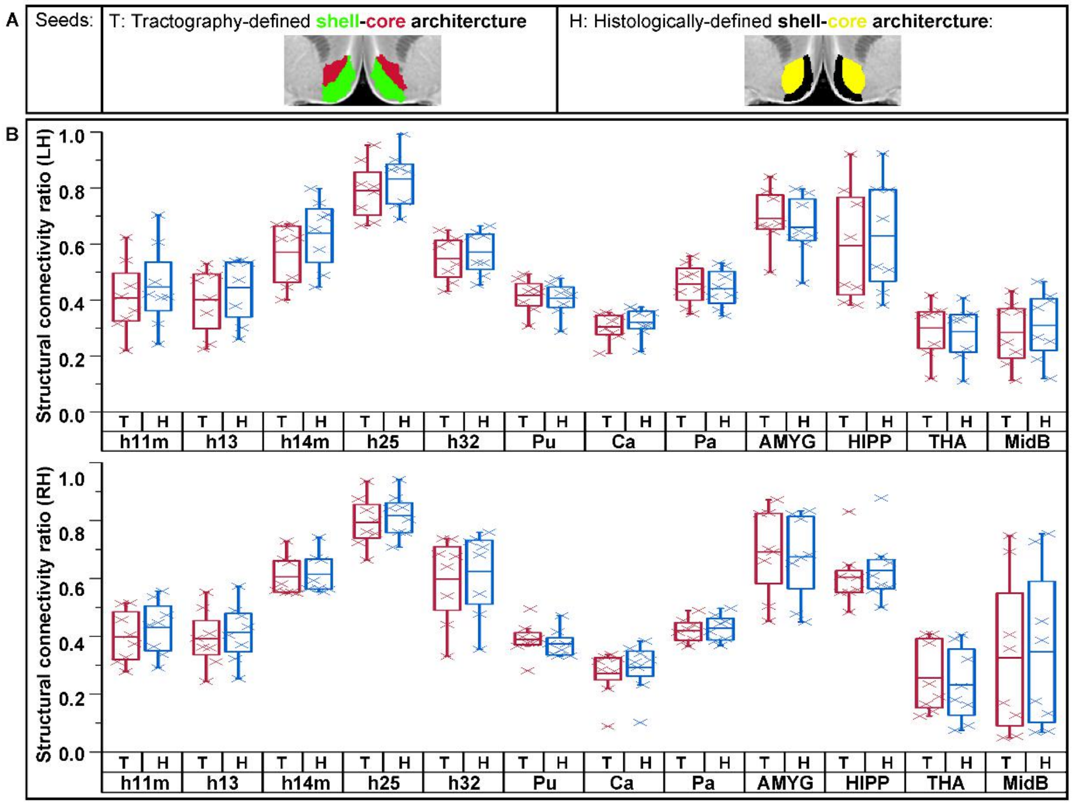


**Figure S12.** Re-do the calculation of structural connectivity of the Acb shell-core architecture. (A) ‘T’ and ‘H’ indicate the tractography- and histologically-defined seeds of the Acb shell-core architecture. (B) Statistical analyses of the single structural connectivity ratio between each target and the two Acb subregions. The ratio groups calculated using tractography- and histologically defined seeds are shown in the red and blue box charts, respectively. No target shows a structural connectivity ratio that differed significantly (at the 5% significance level) from the tractography- and histologically-defined seeds.
